# CRISPRi in *Deinococcus radiodurans*

**DOI:** 10.1101/2022.11.23.517625

**Authors:** Chitra S. Misra, Neha Pandey, Deepti Appukuttan, Devashish Rath

## Abstract

The extremely radiation resistant bacterium, *Deinococcus radiodurans*, is a microbe of importance, both, for studying stress tolerance mechanisms and as a chassis for industrial biotechnology. However, the molecular tools available for use in this organism continue to be limiting. In view of this, the CRISPR-Cas tools provide a large repertoire of applications for gene manipulation. We show the utility of the type I-E Cascade system for knocking down gene expression in this organism. A single-vector system was designed for expression of the Cascade components as well as the crRNA. The type I-E Cascade system was better tolerated than the type II-A Cas9 system in *D. radiodurans*. An assayable acid phosphatase gene, *phoN* integrated into the genome of this organism could be knocked down to 10% of its activity using the Cascade system. Cascade-based knockdown of *ssb*, a gene important for radiation resistance resulted in poor recovery post irradiation. Targeting the Radiation and Desiccation Resistance Motif (RDRM), upstream of the *ssb*, prevented de-repression of its expression upon radiation exposure. In addition to this, multi-locus targeting was demonstrated on the deinococcal genome, by knocking down both *phoN* and *ssb* expression simultaneously. The programmable CRISPRi tool developed in this study will facilitate study of essential genes, hypothetical genes, cis-elements involved in radiation response as well as enable metabolic engineering in this organism. Further the tool is amenable for implementing high-throughput approaches for such studies.

## Introduction

As an extremophile, *Deinococcus radiodurans* (*D. radiodurans*) is well-known for its high tolerance to damage caused by ionizing and UV radiation (Slade & Radman, 2011), (Misra et al., 2013). The bacterium is also extremely resistant to desiccation (Mattimore & Battista, 1996) and the vacuum pressure of space (Kawaguchi et al., 2020) (Ott et al., 2019). Studies over years have attributed various mechanisms including efficient protection of its proteome and efficient repair of damaged DNA to contribute to its survival of extreme conditions (Daly et al., 2010). Apart from serving as a model system to understand extreme stress tolerance, these characteristics have made *D. radiodurans* an attractive organism for biotechnological applications such as bioremediation (Lange et al., 1998) (Brim et al., 2000), production of unique pigments (Jeong et al., 2020) and production of small molecules and metabolites (Li et al., 2019) (Kang et al., 2020).

With the availability of defined promoters, for inducible expression of genes (Lecointe et al., 2004)_(Chen et al., 2019), and selection markers (Smith et al., 1988)(Harris et al., 2004) (Bouthier de la Tour et al., 2009) and the development of shuttle plasmids (Smith et al., 1989) (Masters & Minton, 1992)(Meima & Lidstrom, 2000), recombineering system (Jeong et al., 2017) and conjugation system (Brumwell et al., 2022), the tool-kit for the genetic manipulation of this important organism has been expanding.

Importantly, none of these methods are suitable for targeting essential genes. It is conceivable to use inducible promoter to conditionally complement a chromosomal gene disruption. However, such strategy would not only remove the gene from its chromosomal context but maintaining the native expression level of the target gene would also be a major challenge in the conditional mutant. In most cases these methods are not easy to use, likely due to presence of active restriction-modification systems known to digest exogenous DNA.

Genetic modifications are also difficult to achieve owing to polyploid nature of *D. radiodurans* which can have up to 10 copies of its genome (Hansen, 1978) (Masters et al., 1991). Clustered regularly interspaced short palindromic repeat (CRISPR)-based technology has revolutionized targeted gene manipulation in multitude of organisms (Dominguez et al., 2015). CRISPR editing systems have typically used Cas endonuclease and a paired guide RNA (gRNA). Most gRNAs consist of a fixed sequence derived from the CRISPR repeat and a variable “spacer” region complementary to the targeted locus (Rath et al., 2015a). The gRNA-Cas endonuclease complex interacts with the target locus based on the complementary binding of the spacer region of gRNA to the target. Additionally, this interaction requires presence of a small 3-5 nucleotide long protospacer adjacent motif (PAM) in the target locus. After recognition and binding of the gRNA-Cas endonuclease complex, double-stranded break (DSB) is generated within the target DNA (Rath et al., 2015a). The repair of DSB can proceed via nonhomologous end joining (NHEJ) or via homology-directed repair. For homology-directed repair a donor DNA with homology to the region surrounding the DSB needs to be provided. Both these mechanisms have been exploited for gene editing in many organism (Liu et al., 2022).

CRISPR systems have also been modified to achieve transcriptional repression, a technology referred to as “CRISPRi”. In CRISPR mediated transcriptional repression, mutated version of Cas endonuclease (e.g. dead Cas9 or dCas9) that has lost the nuclease function but retains the ability to bind to the target is used to target the promoter or the ORF region of a gene by use of gRNAs designed for the purpose. Binding of the Cas-gRNA complex to the promoter or the ORF region causes a steric hindrance for the RNA polymerase, thus blocking transcription initiation or elongation (Qi et al., 2013). CRISPR interference was first demonstrated in *Escherichia coli* and mammalian cells (Qi et al., 2013) (Rath et al., 2015) and subsequently has been implemented in a variety of other microbial species (Todor et al., 2021) (Zhang et al., 2021).

As compared to gene editing, CRISPRi has its own advantages. It is relatively easy to engineer for it does not require the presence of a donor template as in the case of editing by homology-directed repair. It allows for regulation of the level of target gene expression with the added advantage that the regulation can be reversed. The CRISPRi can be easily converted to a CRISPR activation (CRISPRa) framework by fusing activator domains to Cas proteins to drive transcriptional activation from a desired locus (Konermann et al., 2014) (Chavez et al., 2015)(Gilbert et al., 2014). Most importantly, it facilitates study of essential genes which cannot be deleted/mutated or for which it is difficult to obtain conditional mutants. So far, CRISPR-based genetic manipulation systems have not been reported in *D. radiodurans*. In this study we report design and development of a type I Cascade based CRISPRi platform for transcription repression of gene expression in *D. radiodurans*.

## Materials and Methods

### Strains and Growth conditions

*Deinococcus radiodurans R1* and *Escherichia coli* JM109 were used in the study. *E. coli* JM109 was used for vector construction and was grown in Luria Bertani (LB) broth. The LB medium was supplemented with appropriate antibiotic (carbenicillin at 100 μg ml^-1^, spectinomycin at 40 μg ml^-1^) wherever required. 1.5 % agar was included in the medium to prepare solid medium. *E. coli* cultures were grown at 37 °C with shaking at 120 rpm. *D. radiodurans* was grown at 32 °C in Tryptone, Glucose, Yeast extract (TGY) with appropriate antibiotics (chloramphenicol 3 μg ml^-1^, spectinomycin 75 μg ml^-1^, kanamycin 8 μg ml^-1^), wherever needed. The amylase activity of the integrated *Deinococcus phoN+* clones was tested on TY starch agar plates as described earlier (Meima et al., 2001). For assaying phosphatase activity on histochemical plates, Phenolphthalein diphosphate (PDP) (1 mg ml^-1^) and methyl green (5 μg ml^-1^) were added to TGY agar. List of strains used in this study is given in Supplementary Table 1.

### Development of CRISPRi platform

The shuttle vector, pRAD1 was used for expression of the CRISPR systems in *D. radiodurans* (Supplementary Table 1). The *cas9* ORF, codon-optimized for *Mycobacterium smegmatis* was analysed for codon usage in *D. radiodurans* using Graphical Codon Analyzer (https://bio.tools/gcua). The *cas9* originally cloned in the pSTKT vector (under communication), was released by digestion with NdeI-BamHI and cloned into identical sites of the pRAD1 vector to generate, pCRD1.

The CRISPR type I-E Cascade operon from *Escherichia coli* K-12 MG1655 was codon optimised for expression in *D. radiodurans* and synthesized along with the sequence coding for the crRNA. The synthesized sequence contained the crRNA sequence under control of the constitutive promoter, P_groESL_ and the ORFs of the codon optimized Cascade subunits. The Shine Dalgarno sequences for each of the Cascade ORFs were retained as it is while optimizing codon usage (Fig. 1a, Supplementary Table 2). The crRNA was designed such that the repeat-spacer-repeat sequence was flanked by SacI-SalI sites for cloning spacers for defining new targets (Fig. 1b). The receiving vector was prepared by first cloning the P_groESL_ in the BamHI-NdeI sites of pRAD1 (Meima & Lidstrom, 2000) to generate pRA-gro. The synthesized Cascade-crRNA fragment was subsequently cloned between the NdeI-XhoI sites of this vector to generate pCRD2. List of plasmids used in the study is given in Supplementary Table 1.

**Fig 1.**
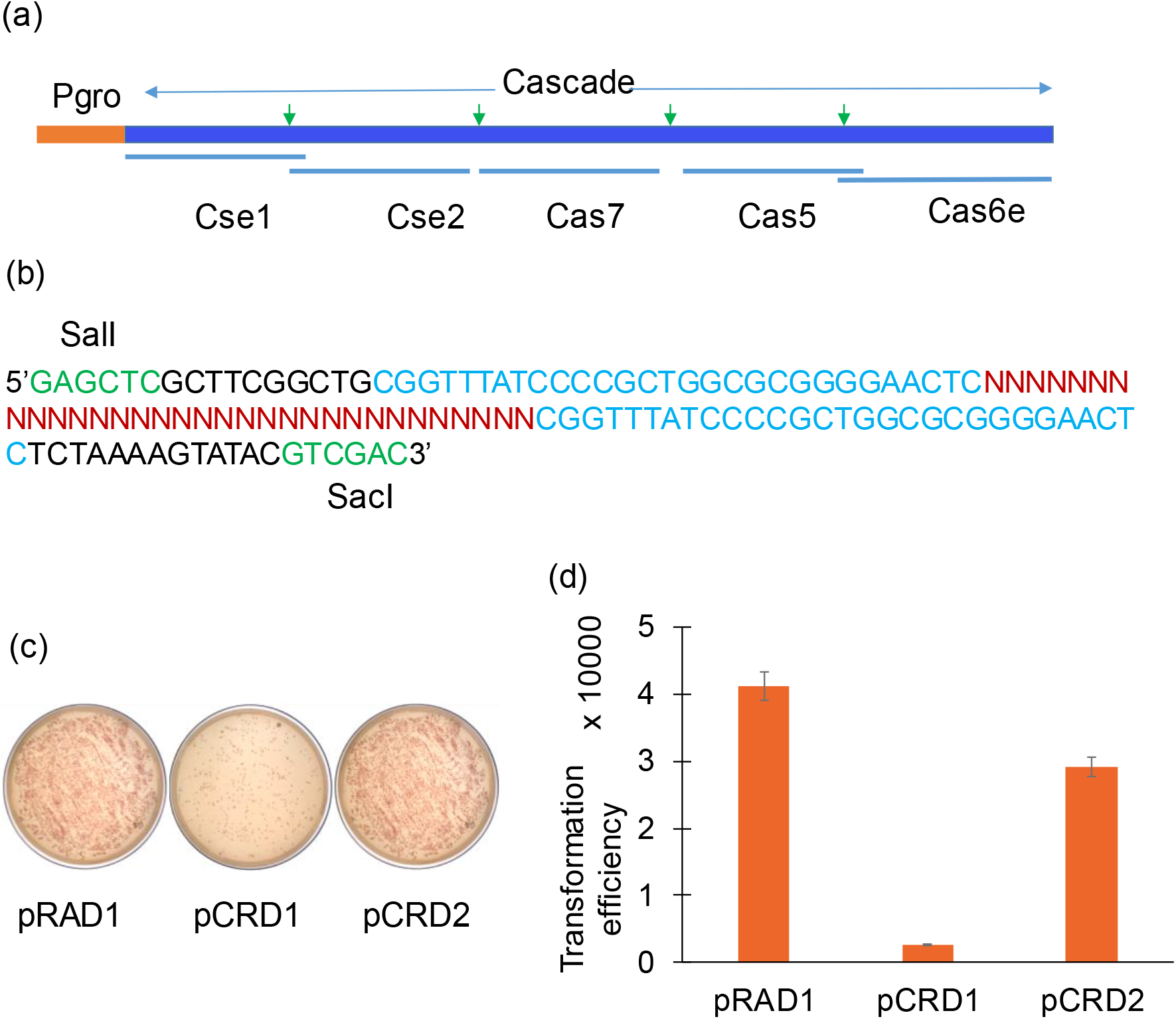
Cascade for gene silencing in *Deinococcus radiodurans*. (a) Codon optimized Cascade operon, with the overlapping nature of ORFs kept intact and Shine Dalgarno (Green arrows) sequences retained. (b) crRNA sequence showing repeat sequences (in blue) on either side of the targeting spacer (in maroon), with flanking restriction sites for cloning marked in green. (c) Plasmids, pRAD1 or pCRD1 or pCRD2 were transformed into *D. radiodurans* and plated on agar plate containing chloramphenicol for selection. (d) Transformation efficiency obtained with various plasmids.

### Integration of *phoN* into the deinococcal chromosome

To generate a stable easy-to-assay system for CRISPRi evaluation in *D. radiodurans*, an acid phosphatase, *phoN*, from *Salmonella typhi* was integrated into chromosome. The recombinant plasmid, pPN1(Appukuttan et al., 2006), which was previously generated in the lab was used as template for amplifying the deinococcal *groESL* promoter along with the *phoN* ORF (P*_groESL_*+*phoN*). The integration plasmid, pGroES4Z, was used as the vector for the transport of the desired insert into the *D. radiodurans* genome (Meima et al., 2001). The plasmid pGroES4Z is an integration vector with a kanamycin cassette flanked by 5’ and 3’ segments of *amyE* gene, which upon integration, replaces the wild type *amyE* gene of *D. radiodurans* leading to loss of starch degrading ability. A 1.2 kb DNA fragment containing *phoN* downstream of deinococcal *groESL* promoter was PCR amplified from plasmid pPN1 using primer pairs P5/P6 and introduced into the XcmI restricted site of pGroES4Z yielding a 9.4 kb recombinant plasmid, pGroES4ZN. A schematic representation of the cloning strategy is shown in Supplementary Figure 2. pGroES4ZN was used to transform *E. coli* DH5α and recombinants were selected on ampicillin plates to obtain *E. coli*-pGroES4ZN clones. The plasmids isolated from these cells were sequenced to rule out mutations and then transformed into *D. radiodurans*. The deinococcal recombinants selected on kanamycin were analysed for integration/recombination event by diagnostic PCR with primers, Amy-1, Amy-2, Amy-3 and Amy-4. The integration was confirmed by DNA sequencing. The details of the primers used are given in Supplementary Table 3.

### Cloning and expression of crRNA targeting different loci on the deinococcal genome

To clone and express sequences coding for the crRNAs for different targets, oligonucleotides of length 90-98 bases were synthesized with overhangs for the SacI and SalI sites. The single stranded oligos (100 μM) were annealed in annealing buffer (10 mM Tris HCL, 50 mM NaCl and 1 mM EDTA) with gradual cooling from 94 °C for 4 min, 75 °C for 5 min, 65 °C for 15 min to 25 °C for 20 min. The annealed oligos were diluted ten times and ligated into pCRD2 digested with SacI and SalI, and transformed into *E. coli* JM109. The transformants were selected on carbenicillin and screened by sequencing of the plasmids to confirm presence of desired target sequence in the crRNA. List of oligonucleotides used for cloning of the crRNA for different targets are listed in Supplementary Table 3.

### Phosphatase assays

Phosphatase assays were carried as described before (Appukuttan et al., 2006). For whole cell assays, cell density was equalized and the cells were incubated with the substrate, *p*-nitrophenol phosphate in acetate buffer (100 mM, pH 5.0) at 37 °C for 1 h. The reaction was stopped by adding 0.2 N NaOH. The product formed due to phosphatase action, *p*-nitrophenol, was estimated by recording absorbance at 405 nm. Activity obtained in *D. radiodurans phoN+* strain was normalized against phosphatase activity obtained in WT strain to quantify contribution of *phoN* alone. In addition to this, phosphatase assay was also determined in the gel by zymogram as described earlier (Appukuttan et al; 2006). Briefly, cells were lysed in non-reducing Laemlli’s buffer, and equal amount of protein was loaded onto gels. Post-electrophoretic separation, the gel was washed in water, followed by Tris buffer (100 mM, pH 8), twice. The gel was subsequently developed using nitroblue tetrazolium (NBT)–5-bromo-4-chloro-3-indolylphosphate (BCIP) mix in 100 mM Tris buffer pH 8.0.

### Irradiation and Growth Curve

Cultures of *D. radiodurans* recombinants were grown overnight, re-suspended at final OD_600nm_ of 3.0 and exposed to Co-60 gamma source to accumulate a cumulative dose of 7 kGy. Cells suspended at similar optical density kept outside the irradiator served as control. The irradiated cells and control cells were subsequently washed in TGY and re-suspended in fresh media to achieve a final OD_600nm_ of 0.5. Cultures were grown for 3 h for post-irradiation recovery (PIR) before harvesting them to extract protein. The Ssb levels in the cells was determined by Western blot using anti-Ssb antibody (Ujaoney et al., 2010). For determining growth kinetics, irradiated and control cells were inoculated into fresh media at a starting OD_600nm_ of 0.1 in microtitre plates. The plate was kept at 32 °C, with shaking (200 rpm) in the plate reader (Infinite M Plex, Tecan Instruments) for acquiring absorbance reading at 600 nm at 30 min interval for 36 h.

## Results

### Development of CRISPRi system

Type II-A Cas9 system from *Streptococcus pyogenes* has been widely adapted for CRISPRi as it involves a single protein Cas9 and a cognate sgRNA. The Cas9 ORF that was earlier codon optimized for *Mycobacterium smegmatis* (under communication) was available in the lab. The codon frequency of *M. smegmatis* and *D. radiodurans* is similar owing to similarly rich GC content of their genome. The *cas9* sequence codon optimized for *M. smegmatis* was tested in silico for codon usage in *D. radiodurans* and found suitable (Supplementary Figure 1). This sequence was cloned into pRAD1 under the constitutive promoter, P_groESL_ to construct pCRD1. The transformation efficiency in *D. radiodurans* was about 16 times lower with pCRD1 than pRAD1 indicating a high level of toxicity for Cas9 (Fig. 1c and 1d). Cas9 is known to cause toxicity in several microbes where it has been applied (Cho et al., 2018), (Rock et al., 2017), (Wendt et al., 2016), (Jiang et al., 2014). In view of the poor TE obtained with the pCRD1 plasmid, we chose to implement the type I-E, Cascade-based system from *E. coli* in *D. radiodurans*. The type I-E system consists of Cas1 and Cas2 (involved in adaptation), a complex of five Cas proteins (Cse1, Cse2, Cas7, Cas5 and Cas6e) named ‘Cascade’ which along with crRNA forms the surveillance complex(Brouns et al., 2008), and Cas3 nuclease which is recruited by Cascade after target recognition for cleavage of the target DNA. For implementing CRISPRi in *D*. *radiodurans*, we designed pCRD2, a single vector system that contained both crRNA expression cassette and an operon coding for Cascade proteins (Fig. 1a and 1b) under control of the strong deinococcal constitutive promoter, P_groESL_. In *D. radiodurans*, the transformation efficiency with pCRD2 was only 1.4 times lesser than the parental vector pRAD1 (Fig. 1c and 1d) indicating better tolerance of the Cascade system in *D. radiodurans* than the Cas9 system.

### Integration of *phoN* into the deinococcal chromosome

To demonstrate that the developed CRISPRi platform can be used for transcriptional repression, *phoN*, which provided an easily assayable phenotype was integrated in the genome of the *D. radiodurans*. For this, pGroES4ZN (a *phoN* carrying derivative of integrative plasmid pGroES4Z) was transformed into *D. radiodurans* and kanamycin-resistant clones obtained were screened for phosphatase activity by plating on media containing PDP-MG. Green colored colonies obtained with the recombinant strain, indicated phosphatase activity as compared to the typical orange colored colony obtained with wild type cells (Fig. 2a). Whole cell *p*NPP phosphatase assay also confirmed the presence of the *phoN* activity (Supplementary Table. 4). The vector pGroES4Z integrates into the *Deinococcus* genome at the *amyE* locus and hence during a double recombination event, there is disruption of this gene, resulting in inability to degrade starch. The wild type *D. radiodurans* and the integrated *Deinococcus phoN*+ clones were analysed for their starch degrading capability on a starch agar plate. A typical result is shown in Fig 2b. Wild type *D. radiodurans* showed a halo zone around its growth on the starch agar plate, when flooded with iodine solution. Under similar conditions, no halo zone was observed with the *Deinococcus phoN+* clone, indicating a disruption in the *amyE* gene. This analysis confirmed integration of the *phoN* in the *amyE* locus of the *D. radiodurans* genome.

**Fig 2.**
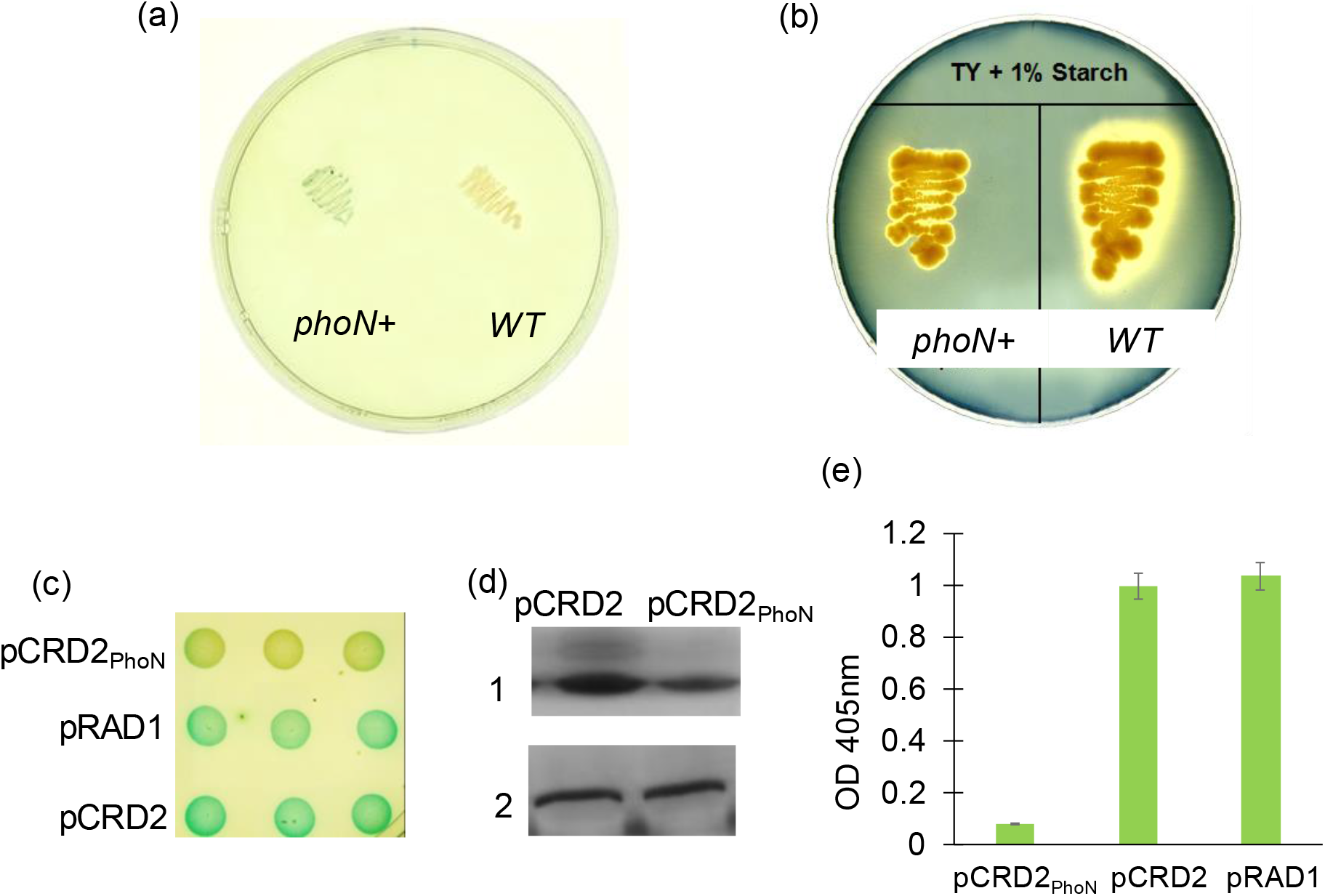
Phosphatase silencing using Cascade in *D. radiodurans phoN*+. The *phoN* gene was integrated into the wild type (WT) *D. radiodurans* R1 cells. Integration of the *phoN* gene into deinococcal chrmososme was evident by (a) PhoN activity as observed on PDP-MG histochemical plate and (b) loss of starch hydrolysis observed on starch agar plate. Phosphatase silencing was assayed by (c) histochemical assays, (d) zymogram and (e) biochemical assay. In the zymogram, 1 refers to the band developed due to PhoN activity and 2 refers to activity of a native phosphatase. Phosphatase activity in *phoN+* cells was normalized against activity in *WT* cells.

### CRISPRi for down-regulation of *phoN* gene

The crRNA was designed to target the sequence immediately downstream of the *phoN* start codon on the chromosome. The sequence coding for the *phoN*-targeting crRNA was cloned in pCRD2 to generate, pCRD2_PhoN_. The plasmid was transformed into *D. radiodurans phoN*+strain and phosphatase activity was screened in the transformants. Cascade mediated silencing of *phoN* was assessed using equal cell densities on histochemical plates where colonies of recombinants carrying pCRD2PhoN showed no green coloration compared to those carrying pCRD2 or pRAD1 that showed intense green color, indicative of *phoN* silencing (Fig. 2c). Likewise, zymogram for phosphatase expression showed presence of a 27 kDa band in protein extracts from cells carrying pCRD2PhoN that was of lesser intensity than a similar sized band in cells carrying pCRD2 further confirming the silencing of the gene (Fig. 2d). Densitometry of the bands showed nine times lower phosphatase activity in protein extracted from pCRD2PhoN compared to pCRD2. On the other hand, a native phosphatase that gave a band of approximately 120 kDa in zymogram, showed equal intensity in protein extracts from both pCRD2_PhoN_ and pCRD2 carrying recombinants and served as a loading control (Fig. 2d). The test and control strains were assessed for *phoN* expression in quantitative phosphatase assays and a 12-fold silencing efficiency was calculated (Fig. 2e).

### CRISPRi to down-regulate *ssb*, a gene involved in radiation resistance

The PhoN assay demonstrated that Cascade-based CRISPRi was functional in *D. radiodurans*. In order to establish it as a general gene silencing tool and to assess physiological relevance of deinococcal gene silencing we chose to knock down a native gene, *ssb* which plays an important role in radiation resistance. As in case of *phoN*, a spacer (SpORF) was targeted at the starting of the *ssb* ORF and cloned into pCRD2 (Fig. 3a) to generate, pCRD2_SpORF_. *D. radiodurans* recombinants carrying pCRD2_SpORF_ gave rise to smaller colonies compared to those carrying pCRD2 (Fig. 3b). The expression of Ssb was analysed on Western blot (Fig. 3c). Control cells carrying pCRD2 showed a basal level of expression. Upon irradiation, the Ssb levels showed a marked increase. Compared to the control cells, cells with pCRD2_SpORF_, showed much lower Ssb levels under unirradiated conditions confirming the knockdown of *ssb* expression. Upon irradiation, in these cells, Ssb levels did increase, but remained lower than irradiated control cells (Fig. 3c). Further, under both control and irradiated conditions, cells containing pCRD2_SpORF_ showed poor growth compared to control cells (Fig. 3d), indicating effective knockdown of Ssb at physiologically relevant levels.

**Fig 3.**
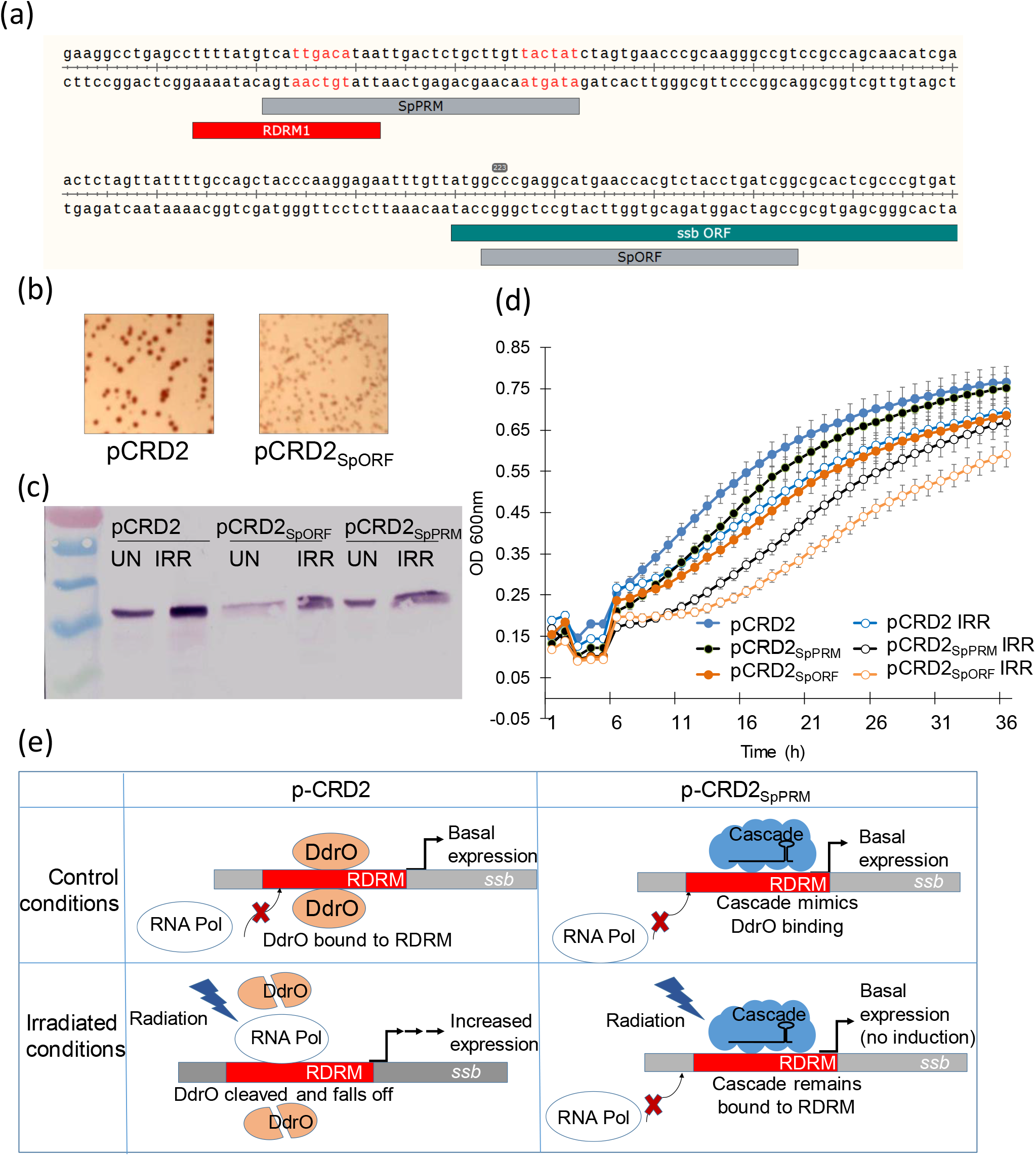
Ssb silencing at promoter and ORF using CRISPR-Cascade system. Two loci, one targeting the promoter (SpPRM) and the other targeting the ORF (SpORF) of the *ssb* gene were identified. The −10 and −35 is marked in red alphabets. The illustration was made using Snapgene software. (b) Smaller sized colonies obtained on plating cells bearing pCRD2SpORF compared to those bearing pCRD2. (c) Western blot for detecting the Ssb protein in total protein extracted from irradiated (IRR) and unirradiated (UN) cells. (d) Growth kinetics during recovery of recombinant cells post-irradiation. (e) Mechanism of induction of *ssb* on irradiation and its repression due to the binding of Cascade to RDRM.

To investigate the effect of Cascade binding to a regulatory sequence element involved in radiation resistance, a spacer (SpPRM) was designed that targeted the promoter of the *ssb* while also overlapping with a Radiation and Desiccation Responsive Motif (RDRM) (Fig. 3a). In cells carrying pCRD2SpPRM, under unirradiated conditions, Ssb level was comparable to unirradiated control cells. However, in the same cells upon irradiation, the Ssb levels barely rose above the basal expression (Fig. 3c). Likewise, such cells showed normal growth under unirradiated conditions, but poor recovery from radiation (Fig. 3d).

Normally, DdrO, a repressor of *ssb* expression remains bound to the RDRM sequences, keeping the Ssb expression to a basal level (Devigne et al., 2015). Upon irradiation, the DdrO is cleaved, relieving repression of *ssb* and inducing its expression(Ludanyi et al., 2014). Our results indicate that crRNA-guided binding of Cascade to RDRM prevents de-repression of the *ssb* as illustrated (Fig. 3e).

### Multiplexed gene silencing in *D. radiodurans*

Ability to target more than one gene simultaneously is one of the main advantages of the CRISPR system. To extend this feature of multiplexed silencing into *D. radiodurans*, a sequence coding for crRNAs targeting the *phoN* and *ssb* was synthesized (Figure 4a) and cloned into the pCRD2 vector to generate pCRD2_SpPhSb_. Recombinant deinococcal cells carrying pCRD2_SpPhSb_, showed lower phosphatase activities, as evidenced by lack of green coloration of colonies on histochemical plate, in contrast to dark green colonies of cells carrying pCRD2 (Figure 4b). pCRD2_SpPhSb_ carrying cells also showed lower levels of Ssb compared to control cells indicating its knockdown (Figure 4c). The silencing of *ssb* was further indicated by poor growth and poor recovery from radiation in growth kinetics experiments (Figure 4d). Together these results showed that two genes could be simultaneously knocked down in *D. radiodurans*.

**Fig 4.**
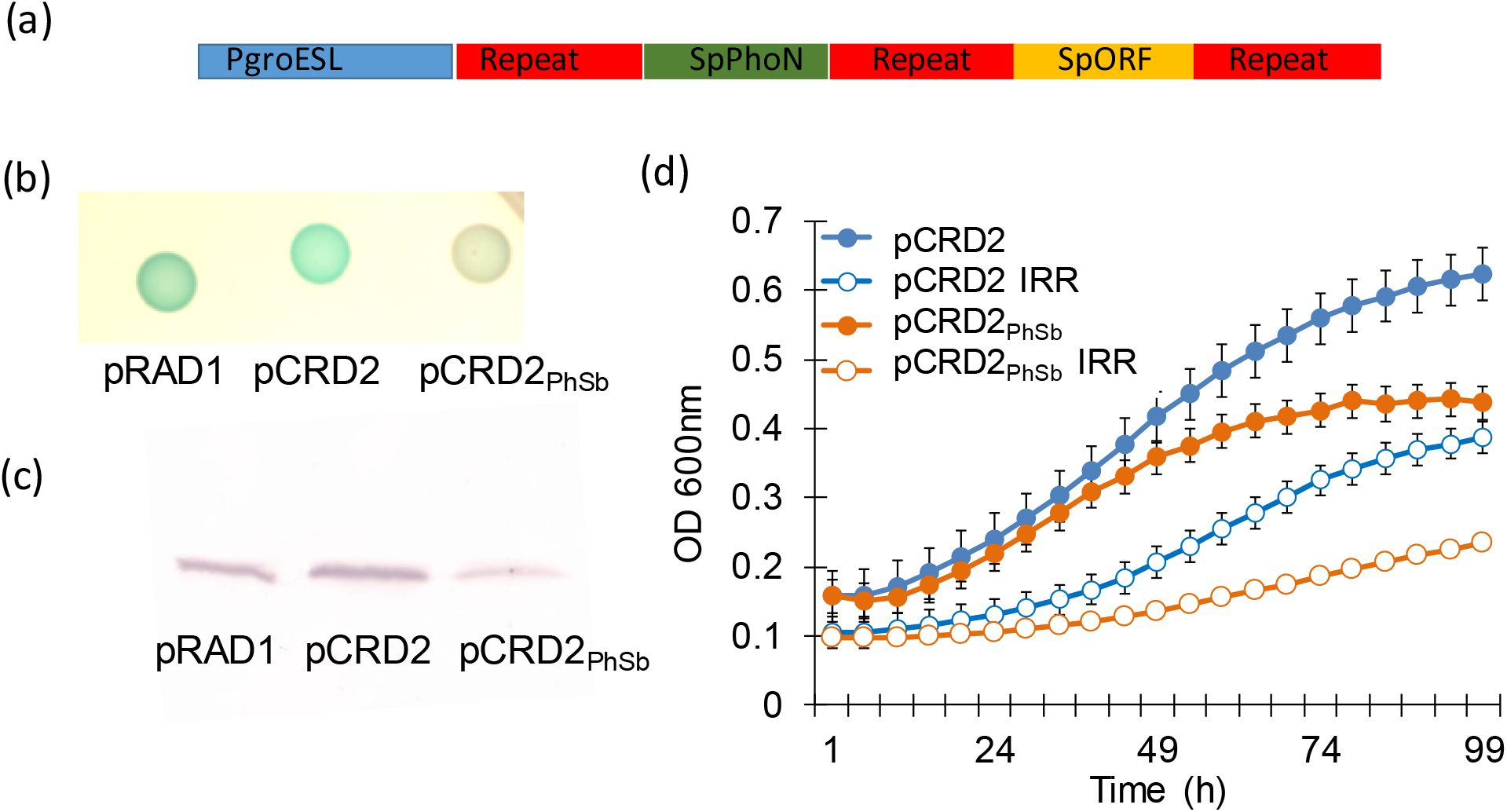
Silencing of *phoN* and *ssb* genes simultaneously in *D. radiodurans* using CRISPR-Cascade system. (a) The cassette expressing the pre-crRNA in the pCRD2SpPhSb plasmid, for targeting *phoN* and *ssb*. (b) Phosphatase silencing as observed in recombinants on histochemical plates. (c) Western blot for detecting the Ssb in total protein extracted from recombinant cells. (d) Growth kinetics of recombinants under irradiated (IRR) and non-irradiated conditions.

## Discussion

*D. radiodurans*, an organism of immense interest to researchers, did not have a system for carrying out targeted gene regulation. Further, multiple copies of the genome and a limited molecular toolkit made it difficult to study the organism. Recently, some systems for targeted gene regulation such as RNA interference and engineered transcription activator-like effector proteins and interference by clustered regularly interspaced short palindromic repeat (CRISPR) sequences have been exploited for regulation of expression in many organisms (Gilbert et al., 2014) (la Russa & Qi, 2015) (Rock et al., 2017) (Tan et al., 2018). While RNA interference is restricted to particular organisms, custom DNA-binding proteins are difficult to implement because of high cost associated with their designing and testing (Rath et al., 2015a). Contrary to these, CRISPR interference (CRISPRi) approach offers a simple and cost-effective tool principally applicable to all microorganisms for targeted gene regulation (Barrangou & Doudna, 2016) (Dominguez et al., 2015).

The dearth of molecular tools in *D. radiodurans* has limited the progress in studying its radiation resistance mechanisms. For example, in well-worked out system of DdrO-based regulation of radiation response, it is still not known how PprI is activated (Lu & Hua, 2021). Not all the transcriptional regulators have been characterized in this microbe and the role of sRNAs in radiation response requires further investigation. Crosstalk among intricate regulatory networks that may play an important role in radiation resistance have not been well-studied owing lack of appropriate tools to regulate expression of multiple genes in this microbe (Wang et al., 2019). The realization of full potential of this organism in industrial biotechnology has also been hampered by absence of convenient genetic tools. In view of this, we report the addition of CRISPR-based gene silencing to the molecular toolkit available for this organism.

We used the type I-E system, as the type II-A Cas9 system was poorly tolerated in this organism, and problems with use of the latter in microbes have earlier been reported due to Cas9 toxicity (Rock et al., 2017) (Wendt et al., 2016). This study demonstrates application of Cascade-based system to knockdown an assayable gene, *phoN* as well as an essential gene, *ssb*. As promoters in this organism are not very well characterized and are poorly predicted by bioinformatics tools (Chen et al., 2019), we targeted the start of the ORF regions to ascertain silencing efficiency, a feature that will have to be necessarily used for functional studies of hypothetical, novel or poorly characterized genes. This is especially important in this organism considering the inability of in silico approaches to efficiently predict even strong deinococcal promoters (Chen et al., 2019). The efficiency of knockdown of *phoN* as assayed by phosphatase activity was around 90%, despite the multipartite genome in this organism.

Similarly, targeting the ORF of *ssb* resulted in low Ssb levels which in turn caused growth defect under both irradiated and unirradiated conditions. Typically, modified CRISPR systems used for silencing of gene expression work best on targeting the promoter. Curiously, targeting the stretch of promoter which overlapped the RDRM sequence of *ssb* did not cause lower Ssb levels and likewise did not affect growth. The RDRM is normally bound by the DdrO, which regulates the expression of several genes upon irradiation. Binding by DdrO keeps the *ssb* expression at a basal level by limiting access of RNA polymerase to the promoter (Devigne et al., 2015). Upon irradiation, DdrO is cleaved by PprI, in turn causing de-repression of *ssb* expression (Ludanyi et al., 2014) (Ujaoney et al., 2010). Binding of Cascade to RDRM repressed *ssb*, mimicking DdrO and resulting in Ssb levels similar to control under unirradiated conditions (Fig. 3e). However, upon irradiation, Cascade binding could not be de-repressed not permitting Ssb induction. This showed the ability of the CRISPR-based gene regulation to characterize cis elements involved in regulation of radiation response, that could also be extended to promoter characterizations.

With Cascade-based CRISPRi we demonstrate the ability to silence two genes simultaneously. This could be expanded to target more genes by engineering the cassette expressing the crRNAs (Li et al., 2015) (Lv et al., 2015). This is an added advantage with Cascade as multi-gene regulation is relatively difficult to achieve with Cas9 systems. Multiplexed gene regulation in *D. radiodurans* will enable interrogation of several genes in a pathway or genes in different interacting pathways. This will aid investigation of various multi-gene phenomena such as radiation resistance apart from finding use in manipulating substrate/product flux for metabolic engineering applications. In addition, CRISPRi screens will provide a high throughput method to probe gene networks. We anticipate that this easy-to-use gene silencing tool will facilitate study of several interesting phenomena such as role of sRNA and unique regulatory pathways in radiation resistance which have not been investigated in-depth in this organism.

## Acknowledgements

Authors thank Dr. AVSSN Rao, Head, Applied Genomics Section, BARC for critical reading of the manuscript. Authors acknowledge Dr. Rita Mukhopadhyaya for her suggestions on applications of this work. Authors thank Dr. Aman Kumar Ujaoney for providing the Anti-Ssb antibody and Shaikh M.I. for the technical help in conduct of experiments.

## References

Appukuttan, D., Rao, A. S., & Apte, S. K. (2006). Engineering of Deinococcus radiodurans R1 for bioprecipitation of uranium from dilute nuclear waste. Applied and Environmental Microbiology, 72(12), 7873–7878. https://doi.org/10.1128/AEM.01362-06

Barrangou, R., & Doudna, J. A. (2016). Applications of CRISPR technologies in research and beyond. Nature Biotechnology, 34(9), 933–941. https://doi.org/10.1038/nbt.3659

Bouthier de la Tour, C., Toueille, M., Jolivet, E., Nguyen, H. H., Servant, P., Vannier, F., & Sommer, S. (2009). The Deinococcus radiodurans SMC protein is dispensable for cell viability yet plays a role in DNA folding. Extremophiles: Life under Extreme Conditions, 13(5), 827–837. https://doi.org/10.1007/S00792-009-0270-2

Brim, H., McFarlan, S. C., Fredrickson, J. K., Minton, K. W., Zhai, M., Wackett, L. P., & Daly, M. J. (2000). Engineering Deinococcus radiodurans for metal remediation in radioactive mixed waste environments. Nature Biotechnology 2000 18:1, 18(1), 85–90. https://doi.org/10.1038/71986

Brouns, S. J. J., Jore, M. M., Lundgren, M., Westra, E. R., Slijkhuis, R. J. H., Snijders, A. P. L., Dickman, M. J., Makarova, K. S., Koonin, E. v., & van der Oost, J. (2008). Small CRISPR RNAs guide antiviral defense in prokaryotes. Science, 321(5891). https://doi.org/10.1126/science.1159689

Brumwell, S. L., van Belois, K. D., Giguere, D. J., Edgell, D. R., & Karas, B. J. (2022). Conjugation-Based Genome Engineering in Deinococcus radiodurans. ACS Synthetic Biology, 11(3), 1068–1076. https://doi.org/10.1021/ACSSYNBIO.1C00524

Chavez, A., Scheiman, J., Vora, S., Pruitt, B. W., Tuttle, M., P R Iyer, E., Lin, S., Kiani, S., Guzman, C. D., Wiegand, D. J., Ter-Ovanesyan, D., Braff, J. L., Davidsohn, N., Housden, B. E., Perrimon, N., Weiss, R., Aach, J., Collins, J. J., & Church, G. M. (2015). Highly efficient Cas9-mediated transcriptional programming. Nature Methods 2015 12:4, 12(4), 326–328. https://doi.org/10.1038/nmeth.3312

Chen, A., Sherman, M. W., Chu, C., Gonzalez, N., Patel, T., & Contreras, L. M. (2019). Discovery and characterization of native Deinococcus radiodurans promoters for tunable gene expression. Applied and Environmental Microbiology, 85(21). https://doi.org/10.1128/AEM.01356-19

Cho, S., Choe, D., Lee, E., Kim, S. C., Palsson, B., & Cho, B.-K. (2018). High-Level dCas9 Expression Induces Abnormal Cell Morphology in Escherichia coli. ACS Synthetic Biology, 7(4), 1085–1094. https://doi.org/10.1021/acssynbio.7b00462

Daly, M. J., Gaidamakova, E. K., Matrosova, V. Y., Kiang, J. G., Fukumoto, R., Lee, D. Y., Wehr, N. B., Viteri, G. A., Berlett, B. S., & Levine, R. L. (2010). Small-Molecule Antioxidant Proteome-Shields in Deinococcus radiodurans. PLOS ONE, 5(9), e12570. https://doi.org/10.1371/JOURNAL.PONE.0012570

Devigne, A., Ithurbide, S., Bouthier de la Tour, C., Passot, F., Mathieu, M., Sommer, S., & Servant, P. (2015). DdrO is an essential protein that regulates the radiation desiccation response and the apoptotic-like cell death in the radioresistant Deinococcus radiodurans bacterium. Molecular Microbiology, 96(5), 1069–1084. https://doi.org/10.1111/MMI.12991

Dominguez, A. A., Lim, W. A., & Qi, L. S. (2015). Beyond editing: repurposing CRISPR– Cas9 for precision genome regulation and interrogation. Nature Reviews Molecular Cell Biology 2015 17:1, 17(1), 5–15. https://doi.org/10.1038/nrm.2015.2

Gilbert, L. A., Horlbeck, M. A., Adamson, B., Villalta, J. E., Chen, Y., Whitehead, E. H., Guimaraes, C., Panning, B., Ploegh, H. L., Bassik, M. C., Qi, L. S., Kampmann, M., & Weissman, J. S. (2014). Genome-Scale CRISPR-Mediated Control of Gene Repression and Activation. Cell, 159(3), 647–661. https://doi.org/10.1016/J.CELL.2014.09.029

Hansen, M. T. (1978). Multiplicity of genome equivalents in the radiation-resistant bacterium Micrococcus radiodurans. Journal of Bacteriology, 134(1), 71–75. https://doi.org/10.1128/JB.134.1.71-75.1978

Harris, D. R., Tanaka, M., Saveliev, S. v., Jolivet, E., Earl, A. M., Cox, M. M., & Battista, J. R. (2004). Preserving Genome Integrity: The DdrA Protein of Deinococcus radiodurans R1. PLOS Biology, 2(10), e304. https://doi.org/10.1371/JOURNAL.PBIO.0020304

Jeong, S. W., Kim, J. H., Kim, J. W., Kim, C. Y., Kim, S. Y., & Choi, Y. J. (2020). Metabolic Engineering of Extremophilic Bacterium Deinococcus radiodurans for the Production of the Novel Carotenoid Deinoxanthin. Microorganisms 2021, Vol. 9, Page 44, 9(1), 44. https://doi.org/10.3390/MICROORGANISMS9010044

Jeong, S. W., Yang, J. E., Im, S., & Choi, Y. J. (2017). Development of Cre-lox based multiple knockout system in Deinococcus radiodurans R1. Korean Journal of Chemical Engineering 2017 34:6, 34(6), 1728–1733. https://doi.org/10.1007/S11814-017-0082-5

Jiang, W., Brueggeman, A. J., Horken, K. M., Plucinak, T. M., & Weeks, D. P. (2014). Successful transient expression of Cas9 and single guide RNA genes in Chlamydomonas reinhardtii. Eukaryotic Cell, 13(11), 1465–1469. https://doi.org/10.1128/EC.00213-14

Kang, C. K., Jeong, S. W., Yang, J. E., & Choi, Y. J. (2020). High-Yield Production of Lycopene from Corn Steep Liquor and Glycerol Using the Metabolically Engineered Deinococcus radiodurans R1 Strain. Journal of Agricultural and Food Chemistry, 68(18), 5147–5153. https://doi.org/10.1021/ACS.JAFC.0C01024

Kawaguchi, Y., Shibuya, M., Kinoshita, I., Yatabe, J., Narumi, I., Shibata, H., Hayashi, R., Fujiwara, D., Murano, Y., Hashimoto, H., Imai, E., Kodaira, S., Uchihori, Y., Nakagawa, K., Mita, H., Yokobori, S. I., & Yamagishi, A. (2020). DNA Damage and Survival Time Course of Deinococcal Cell Pellets During 3 Years of Exposure to Outer Space. Frontiers in Microbiology, 11, 2050. https://doi.org/10.3389/FMICB.2020.02050/BIBTEX

Konermann, S., Brigham, M. D., Trevino, A. E., Joung, J., Abudayyeh, O. O., Barcena, C., Hsu, P. D., Habib, N., Gootenberg, J. S., Nishimasu, H., Nureki, O., & Zhang, F. (2014). Genome-scale transcriptional activation by an engineered CRISPR-Cas9 complex. Nature 2014 517:7536, 517(7536), 583–588. https://doi.org/10.1038/nature14136

la Russa, M. F., & Qi, L. S. (2015). The New State of the Art: Cas9 for Gene Activation and Repression. Molecular and Cellular Biology, 35(22), 3800–3809. https://doi.org/10.1128/MCB.00512-15

Lange, C. C., Wackett, L. P., Minton, K. W., & Daly, M. J. (1998). Engineering a recombinant Deinococcus radiodurans for organopollutant degradation in radioactive mixed waste environments. Nature Biotechnology 1998 16:10, 16(10), 929–933. https://doi.org/10.1038/nbt1098-929

Larson, M. H., Gilbert, L. A., Wang, X., Lim, W. A., Weissman, J. S., & Qi, L. S. (2013). CRISPR interference (CRISPRi) for sequence-specific control of gene expression. Nature Protocols 2013 8:11, 8(11), 2180–2196. https://doi.org/10.1038/nprot.2013.132

Lecointe, F., Coste, G., Sommer, S., & Bailone, A. (2004). Vectors for regulated gene expression in the radioresistant bacterium Deinococcus radiodurans. Gene, 336(1), 25–35. https://doi.org/10.1016/J.GENE.2004.04.006

Li, J., Webster, T. J., & Tian, B. (2019). Functionalized Nanomaterial Assembling and Biosynthesis Using the Extremophile Deinococcus radiodurans for Multifunctional Applications. Small, 15(20), 1900600. https://doi.org/10.1002/SMLL.201900600

Li, Y., Lin, Z., Huang, C., Zhang, Y., Wang, Z., Tang, Y., Chen, T., & Zhao, X. (2015). Metabolic engineering of Escherichia coli using CRISPR–Cas9 meditated genome editing. Metabolic Engineering, 31, 13–21. https://doi.org/10.1016/J.YMBEN.2015.06.006

Liu, G., Lin, Q., Jin, S., & Gao, C. (2022). The CRISPR-Cas toolbox and gene editing technologies. Molecular Cell, 82(2), 333–347. https://doi.org/10.1016/J.MOLCEL.2021.12.002

Lu, H., & Hua, Y. (2021). PprI: The Key Protein in Response to DNA Damage in Deinococcus. In Frontiers in Cell and Developmental Biology (Vol. 8). https://doi.org/10.3389/fcell.2020.609714

Ludanyi, M., Blanchard, L., Dulermo, R., Brandelet, G., Bellanger, L., Pignol, D., Lemaire, D., & de Groot, A. (2014). Radiation response in Deinococcus deserti: IrrE is a metalloprotease that cleaves repressor protein DdrO. Molecular Microbiology, 94(2), 434–449. https://doi.org/10.1111/MMI.12774

Lv, L., Ren, Y.-L., Chen, J.-C., Wu, Q., & Chen, G.-Q. (2015). Application of CRISPRi for prokaryotic metabolic engineering involving multiple genes, a case study: Controllable P(3HB-co-4HB) biosynthesis. Metabolic Engineering, 29, 160–168. https://doi.org/10.1016/J.YMBEN.2015.03.013

Masters, C. I., & Minton, K. W. (1992). Promoter probe and shuttle plasmids for Deinococcus radiodurans. Plasmid, 28(3), 258–261. https://doi.org/10.1016/0147-619X(92)90057-H

Masters, C. I., Smith, M. D., Gutman, P. D., & Minton, K. W. (1991). Heterozygosity and instability of amplified chromosomal insertions in the radioresistant bacterium Deinococcus radiodurans. Journal of Bacteriology, 173(19), 6110–6117. https://doi.org/10.1128/JB.173.19.6110-6117.1991

Mattimore, V., & Battista, J. R. (1996). Radioresistance of Deinococcus radiodurans: functions necessary to survive ionizing radiation are also necessary to survive prolonged desiccation. Journal of Bacteriology, 178(3), 633–637. https://doi.org/10.1128/JB.178.3.633-637.1996

Meima, R., & Lidstrom, M. E. (2000). Characterization of the minimal replicon of a cryptic Deinococcus radiodurans SARK plasmid and development of versatile Escherichia coli-D. radiodurans shuttle vectors. Applied and Environmental Microbiology, 66(9), 3856–3867. https://doi.org/10.1128/AEM.66.9.3856-3867.2000

Meima, R., Rothfuss, H. M., Gewin, L., & Lidstrom, M. E. (2001). Promoter cloning in the radioresistant bacterium Deinococcus radiodurans. Journal of Bacteriology, 183(10), 3169–3175. https://doi.org/10.1128/JB.183.10.3169-3175.2001

Misra, H., Rajpurohit, Y. S., & Kota, S. (2013). Physiological and molecular basis of extreme radioresistance in Deinococcus radiodurans. Current Science, 104(2), 194–205.

Ott, E., Kawaguchi, Y., Özgen, N., Yamagishi, A., Rabbow, E., Rettberg, P., Weckwerth, W., & Milojevic, T. (2019). Proteomic and Metabolomic Profiling of Deinococcus radiodurans Recovering after Exposure to Simulated Low Earth Orbit Vacuum Conditions. Frontiers in Microbiology, 10(APR), 909. https://doi.org/10.3389/FMICB.2019.00909/BIBTEX

Qi, L. S., Larson, M. H., Gilbert, L. A., Doudna, J. A., Weissman, J. S., Arkin, A. P., & Lim, W. A. (2013). Repurposing CRISPR as an RNA-Guided Platform for Sequence-Specific Control of Gene Expression. Cell, 152(5), 1173–1183. https://doi.org/10.1016/J.CELL.2013.02.022

Rath, D., Amlinger, L., Hoekzema, M., Devulapally, P. R., & Lundgren, M. (2015). Efficient programmable gene silencing by Cascade. Nucleic Acids Research, 43(1), 237–246. https://doi.org/10.1093/nar/gku1257

Rath, D., Amlinger, L., Rath, A., & Lundgren, M. (2015a). The CRISPR-Cas immune system: Biology, mechanisms and applications. Biochimie, 117, 119–128. https://doi.org/10.1016/j.biochi.2015.03.025

Rock, J. M., Hopkins, F. F., Chavez, A., Diallo, M., Chase, M. R., Gerrick, E. R., Pritchard, J. R., Church, G. M., Rubin, E. J., Sassetti, C. M., Schnappinger, D., & Fortune, S. M. (2017). Programmable transcriptional repression in mycobacteria using an orthogonal CRISPR interference platform. Nature Microbiology 2017 2:4, 2(4), 1–9. https://doi.org/10.1038/nmicrobiol.2016.274

Slade, D., & Radman, M. (2011). Oxidative stress resistance in Deinococcus radiodurans. Microbiology and Molecular Biology Reviews: MMBR, 75(1), 133–191. https://doi.org/10.1128/MMBR.00015-10

Smith, M. D., Abrahamson, R., & Minton, K. W. (1989). Shuttle plasmids constructed by the transformation of an Escherichia coli cloning vector into two Deinococcus radiodurans plasmids. Plasmid, 22(2), 132–142. https://doi.org/10.1016/0147-619X(89)90022-X

Smith, M. D., Lennon, E., McNeil, L. B., & Minton, K. W. (1988). Duplication insertion of drug resistance determinants in the radioresistant bacterium Deinococcus radiodurans. Journal of Bacteriology, 170(5), 2126–2135. https://doi.org/10.1128/JB.170.5.2126-2135.1988

Tan, S. Z., Reisch, C. R., & Prather, K. L. J. (2018). A robust CRISPR interference gene repression system in Pseudomonas. Journal of Bacteriology, 200(7). https://doi.org/10.1128/JB.00575-17/

Todor, H., Silvis, M. R., Osadnik, H., & Gross, C. A. (2021). Bacterial CRISPR screens for gene function. Current Opinion in Microbiology, 59, 102–109. https://doi.org/10.1016/J.MIB.2020.11.005

Ujaoney, A. K., Potnis, A. A., Kane, P., Mukhopadhyaya, R., & Apte, S. K. (2010). Radiation desiccation response motif-like sequences are involved in transcriptional activation of the Deinococcal ssb gene by ionizing radiation but not by desiccation. Journal of Bacteriology, 192(21), 5637–5644. https://doi.org/10.1128/JB.00752-10

Wang, W., Ma, Y., He, J., Qi, H., Xiao, F., & He, S. (2019). Gene regulation for the extreme resistance to ionizing radiation of Deinococcus radiodurans. In Gene (Vol. 715). https://doi.org/10.1016/j.gene.2019.144008

Wendt, K. E., Ungerer, J., Cobb, R. E., Zhao, H., & Pakrasi, H. B. (2016). CRISPR/Cas9 mediated targeted mutagenesis of the fast growing cyanobacterium Synechococcus elongatus UTEX 2973. Microbial Cell Factories, 15(1), 115. https://doi.org/10.1186/s12934-016-0514-7

Zhang, R., Xu, W., Shao, S., & Wang, Q. (2021). Gene Silencing Through CRISPR Interference in Bacteria: Current Advances and Future Prospects. Frontiers in Microbiology, 12. https://doi.org/10.3389/FMICB.2021.635227

