## Supplementary for "CRISPRi in *Deinococcus radiodurans*"

### Supplementary Figures

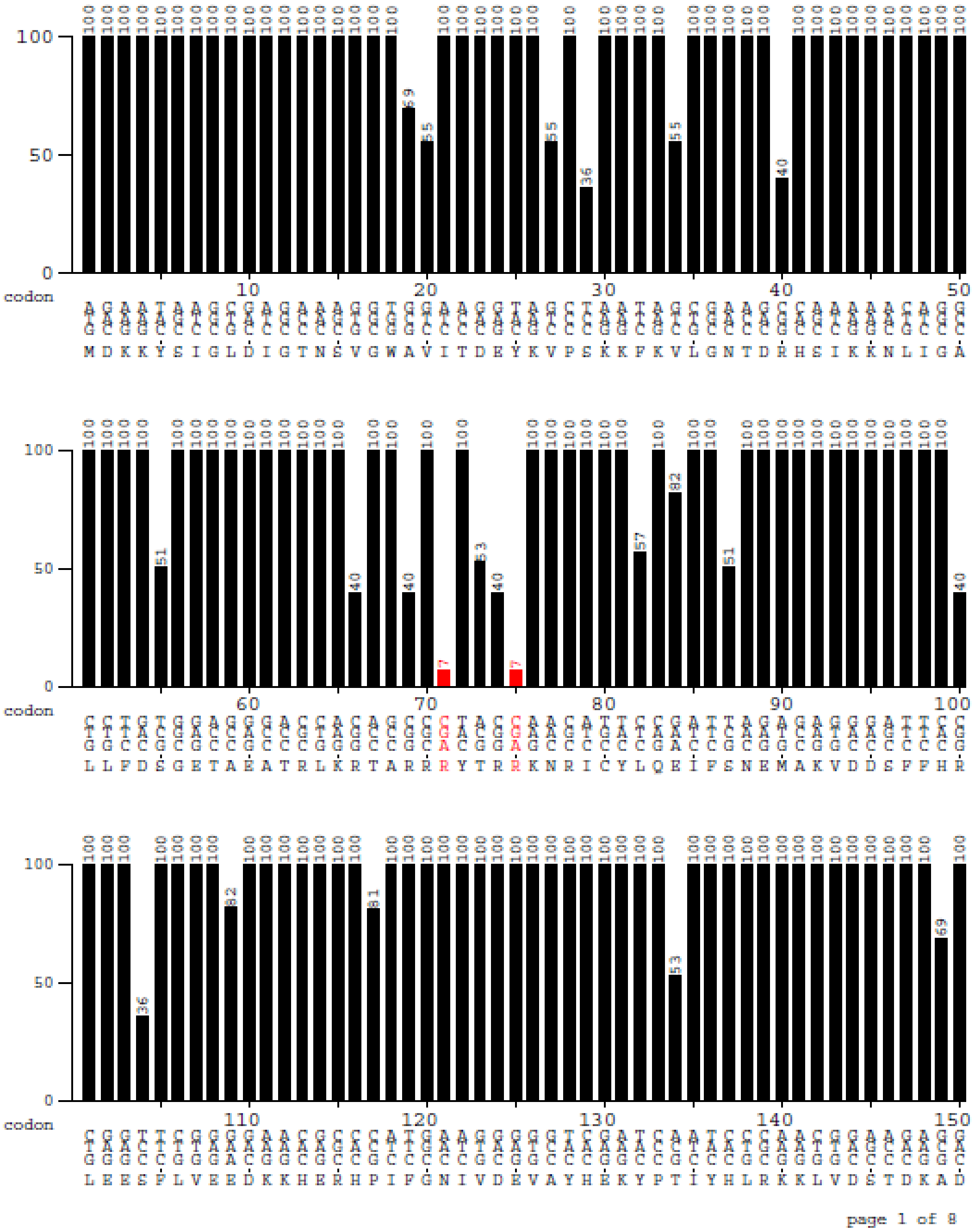

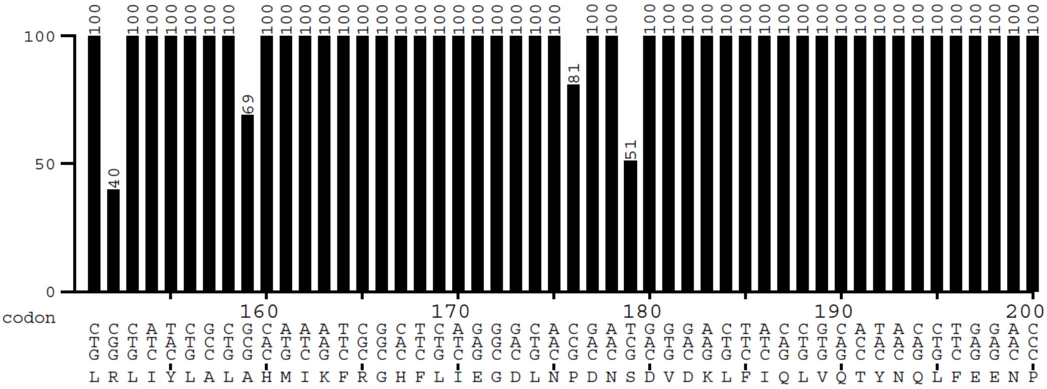
100
50
0
codon
160 170 180 190 200
L R L I Y L A L A H M I K F R G H F L I E G D L N P D N S D V D K L F I Q L V Q T Y N Q L F E E N P

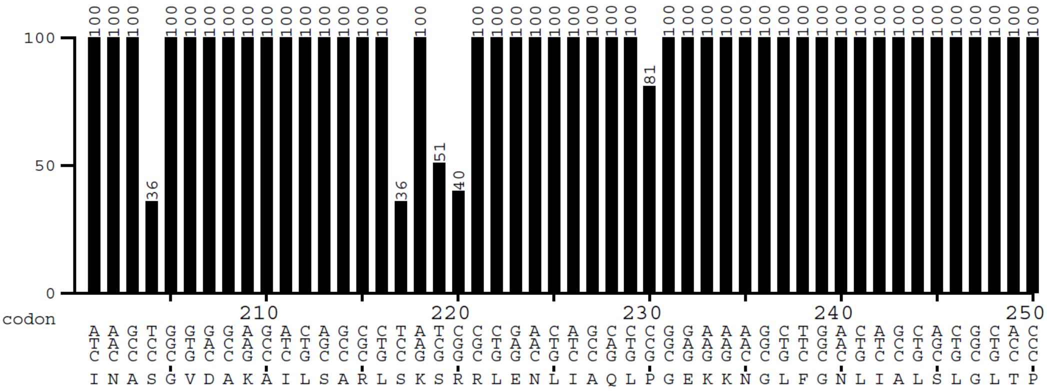
100
50
0
codon
210 220 230 240 250
I N A S G V D A K A I L S A R L S K S R R L E N L I A Q L P G E K K N G L F G N L I A L S L G L T P

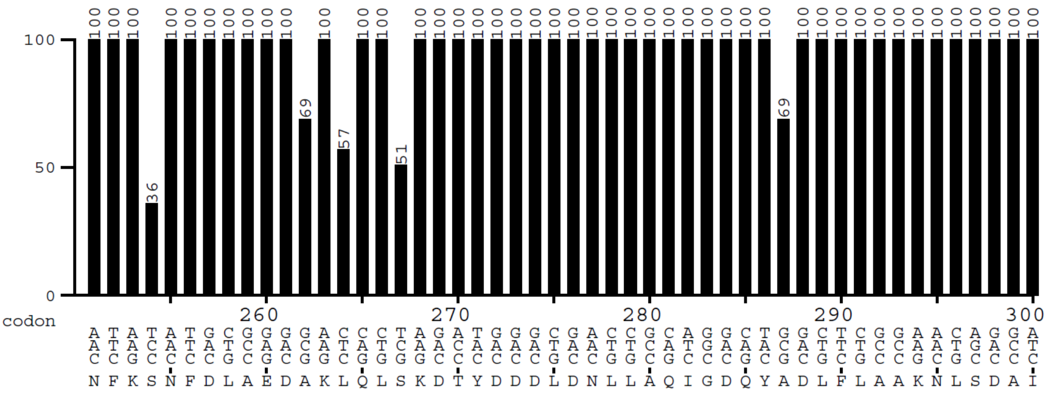
100
50
0
codon
260 270 280 290 300
N F K S N F D L A E D A K L Q L S K D T Y D D D L D N L L A Q I G D Q Y A D L F L A A K N L S D A I

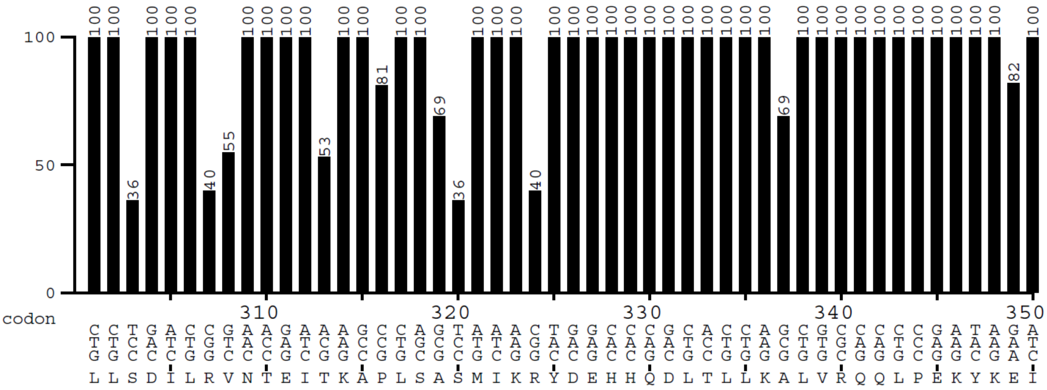
100
50
0
codon
310 320 330 340 350
L L S D I L R V N T E I T K A P L S A S M I K R Y D E H H Q D L T L L K A L V R Q Q L P E K Y K E I

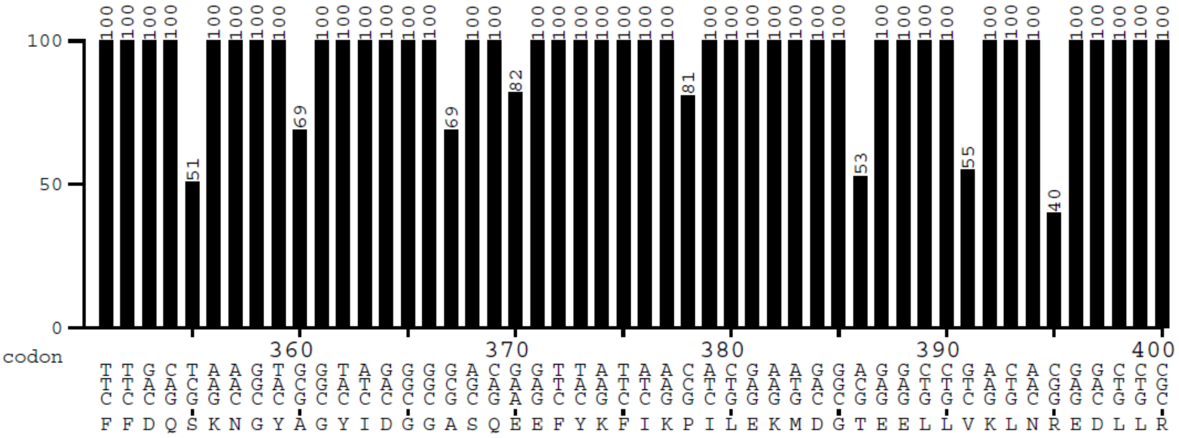
100
50
0
codon
360 370 380 390 400
F F D Q S K N G Y A G Y I D G G A S Q E E F Y K F I K P I L E K M D G T E E L L V K L N R E D L L R

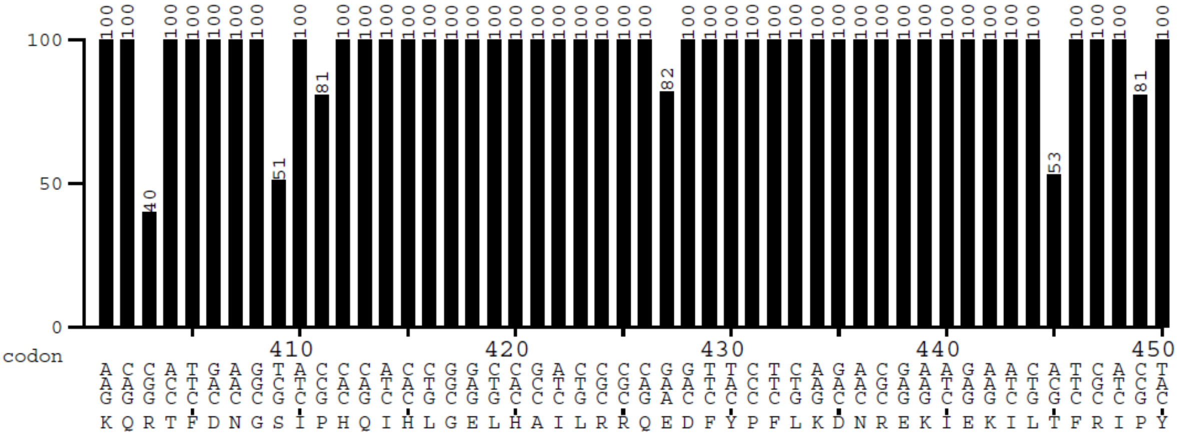
100
50
0
codon
410 420 430 440 450
K Q R T F D N G S I P H Q I H L G E L H A I L R R Q E D F Y P F L K D N R E K I E K I L T F R I P Y

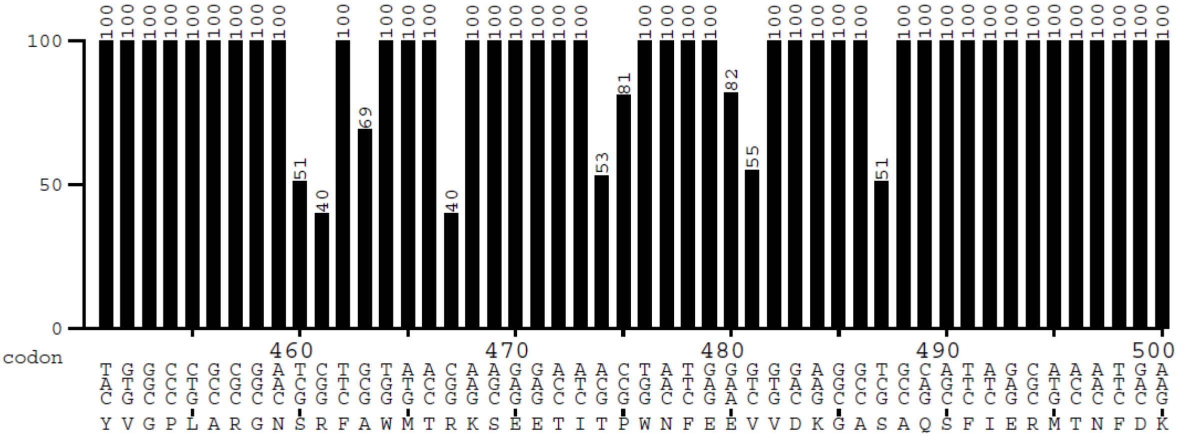
100
50
0
codon
460 470 480 490 500
Y V G P L A R G N S R F A W M T R K S E E T I T P W N F E E V V D K G A S A Q S F I E R M T N F D K

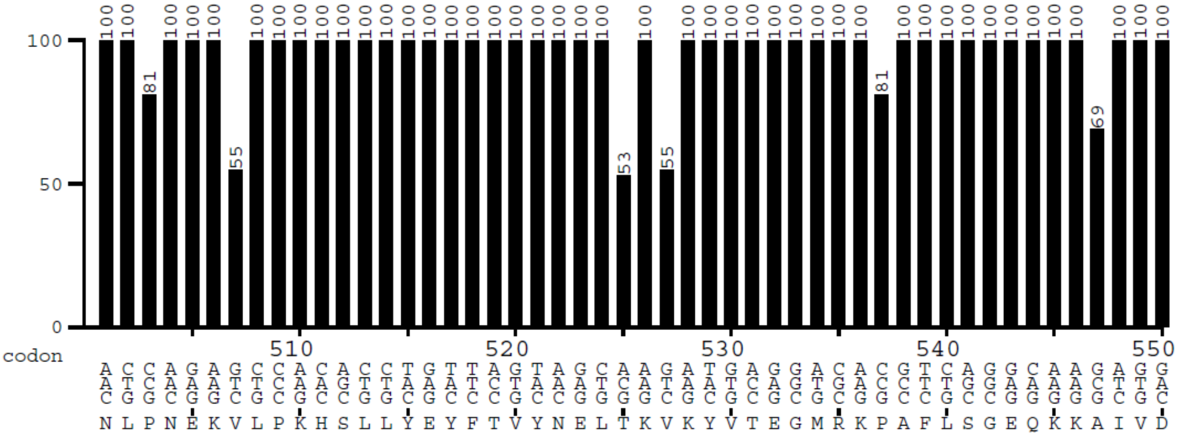
100
50
0
codon
510 520 530 540 550
N L P N E K V L P K H S L L Y E Y F T V Y N E L T K V K Y V T E G M R K P A F L S G E Q K K A I V D

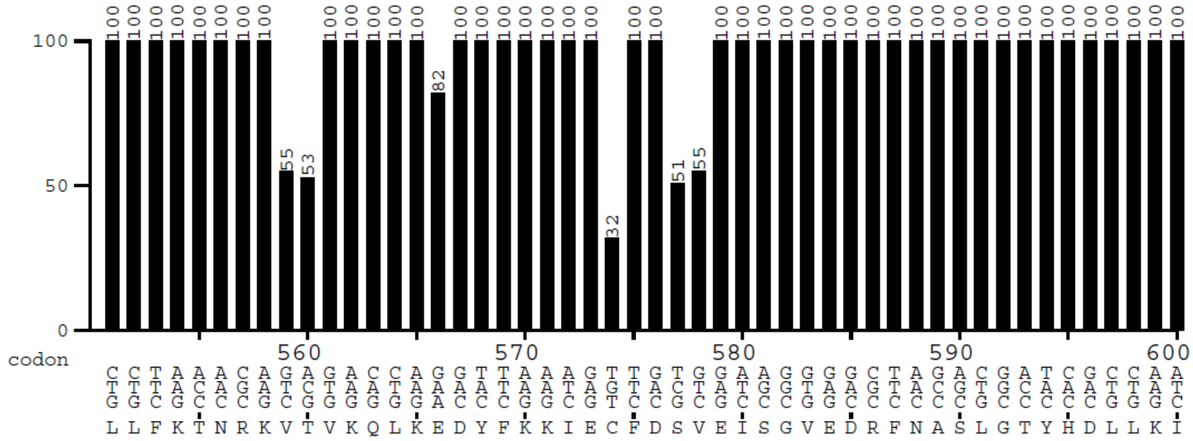
100
50
0
codon
560 570 580 590 600
L L F K T N R K V T V K Q L K E D Y F K K I E C F D S V E I S G V E D R F N A S L G T Y H D L L K I

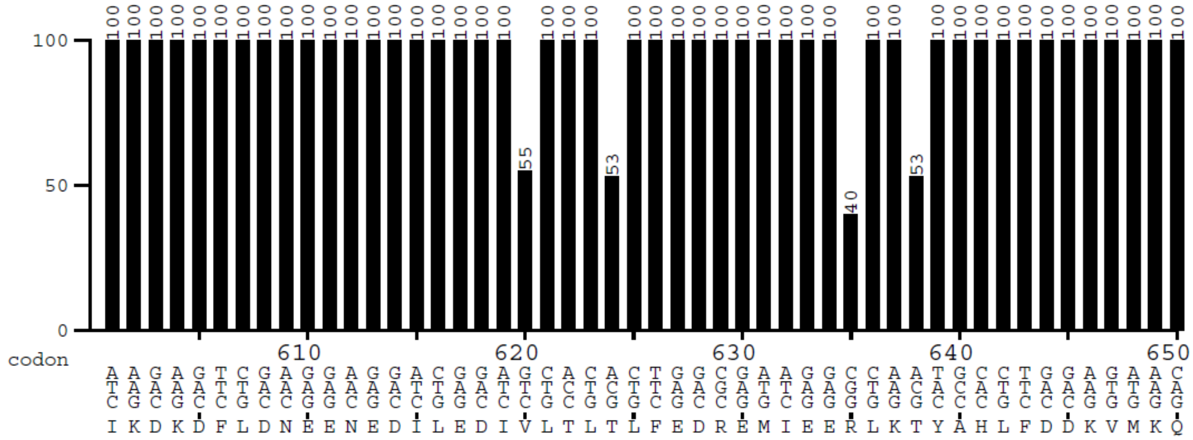
100
50
0
codon
610 620 630 640 650
I K D K D F L D N E E N E D I L E D I V L T L T L F E D R E M I E E R L K T Y A H L F D D K V M K Q

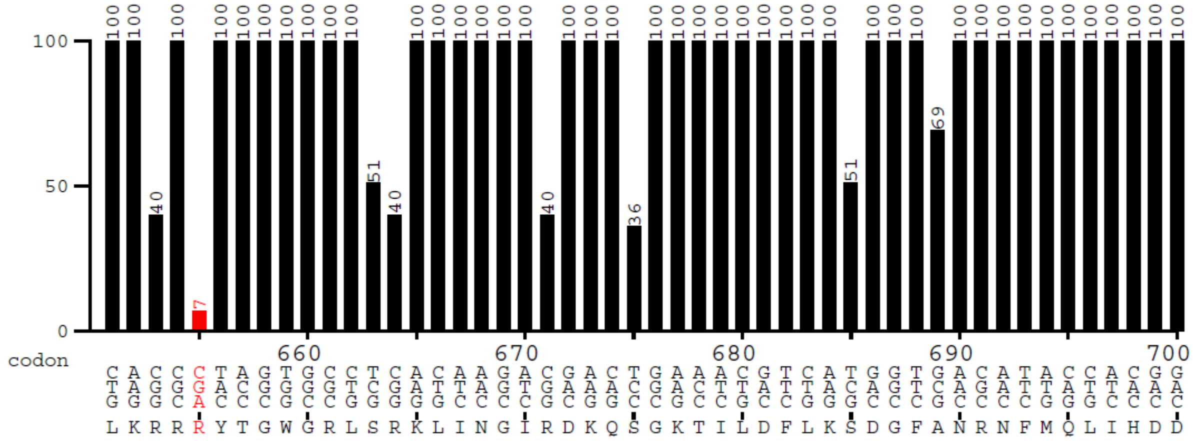
100
50
0
codon
660 670 680 690 700
L K R R R Y T G W G R L S R K L I N G I R D K Q S G K T I L D F L K S D G F A N R N F M Q L I H D D

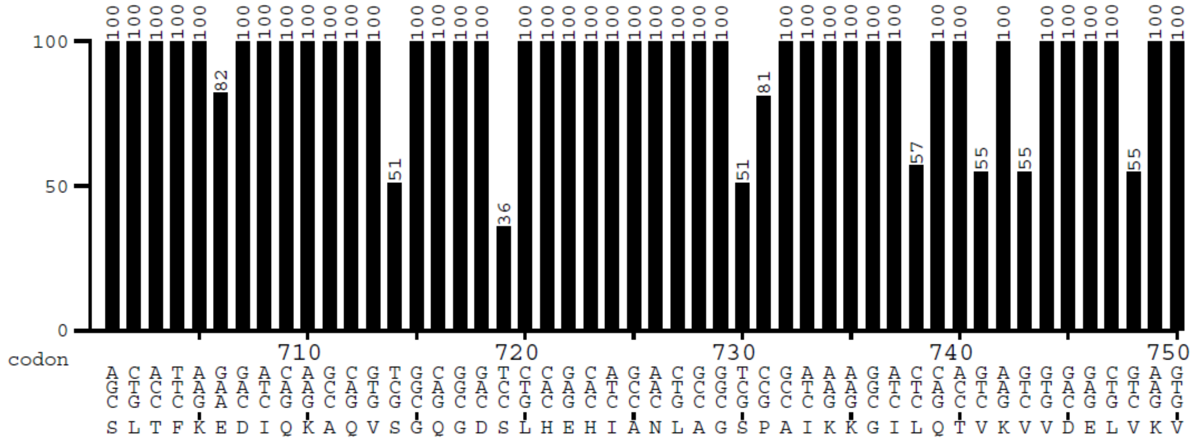
100
50
0
codon
710 720 730 740 750
S L T F K E D I Q K A Q V S G Q G D S L H E H I A N L A G S P A I K K G I L Q T V K V V D E L V K V

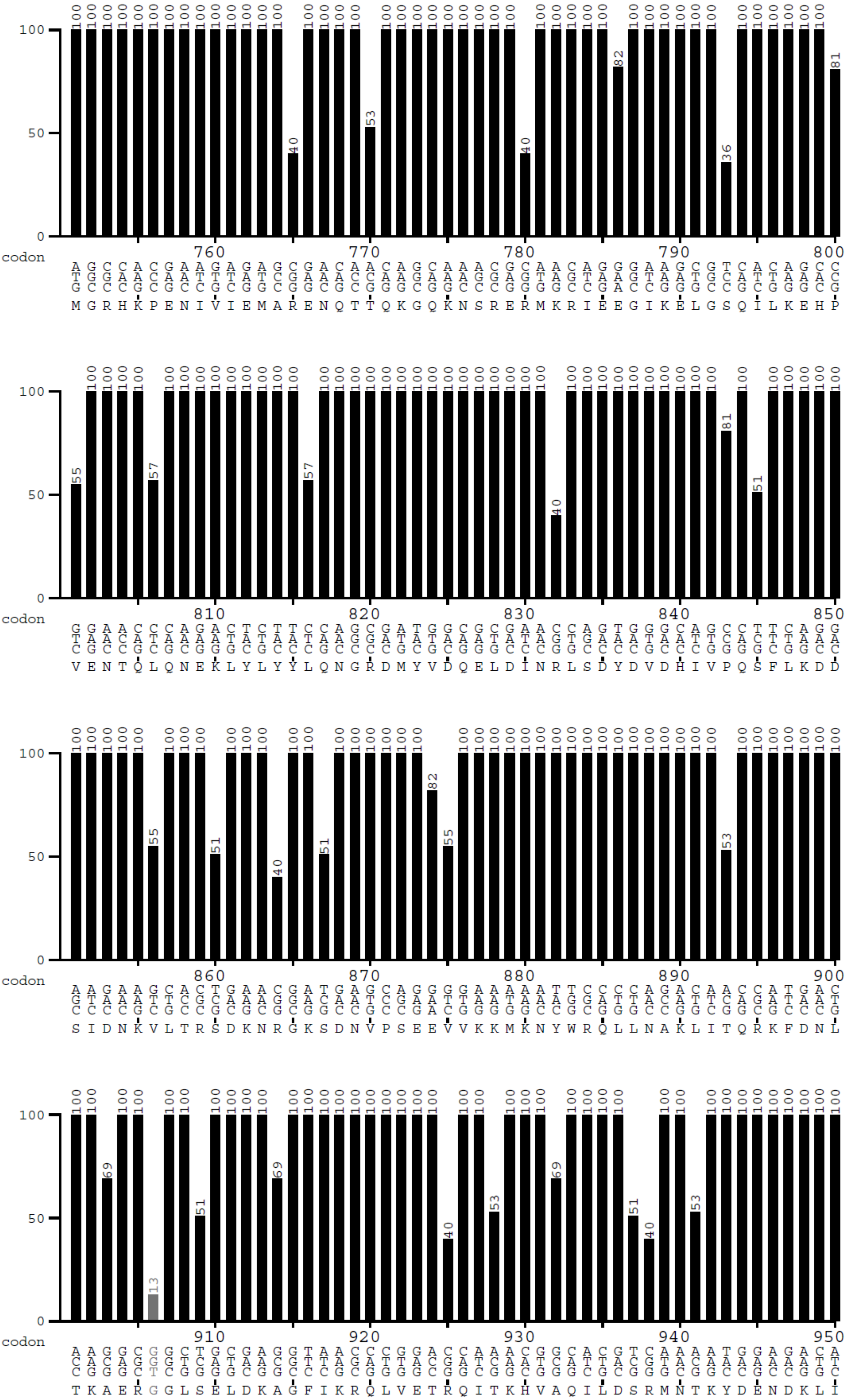
codon
760 770 780 790 800
M G R H K P E N I V I E M A R E N Q T T Q K G Q K N S R E R M K R I E E G I K E L G S Q I L K E H P
810 820 830 840 850
V E N T Q L Q N E K L Y L Y Y L Q N G R D M Y V D Q E L D I N R L S D Y D V D H I V P Q S F L K D D
860 870 880 890 900
S I D N K V L T R S D K N R G K S D N V P S E E V V K K M K N Y W R Q L L N A K L I T Q R K F D N L
910 920 930 940 950
T K A E R G G L S E L D K A G F I K R Q L V E T R Q I T K H V A Q I L D S R M N T K Y D E N D K L I

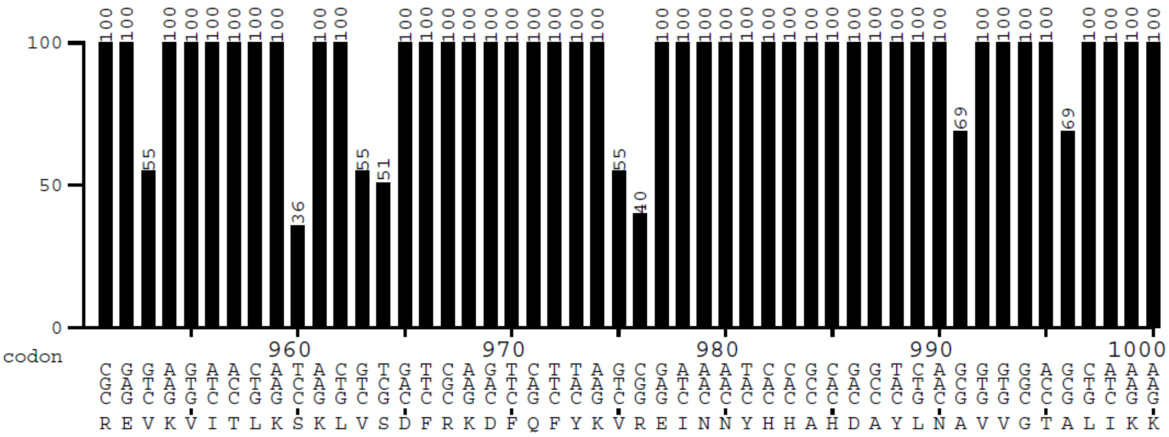
100
50
0
codon
960 970 980 990 1000
R E V K V I T L K S K L V S D F R K D F Q F Y K V R E I N N Y H H A H D A Y L N A V V G T A L I K K

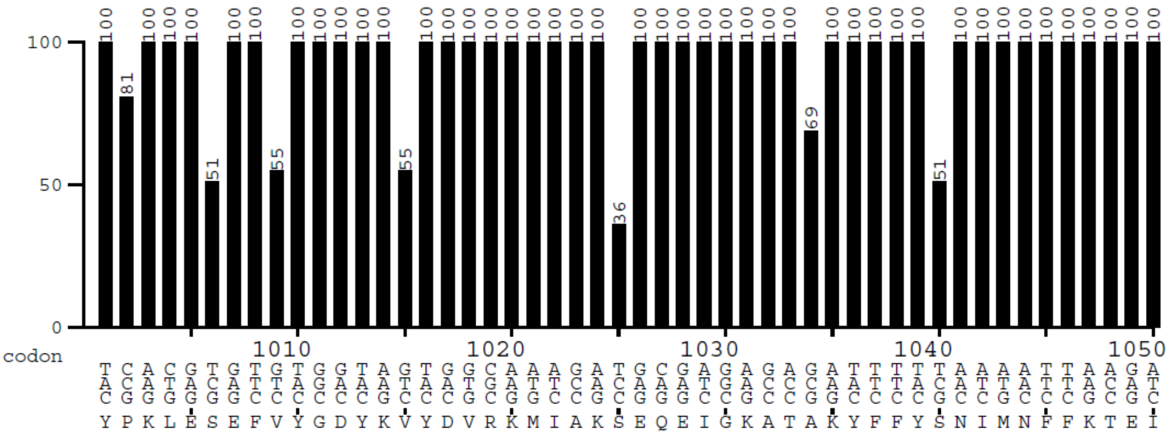
100
50
0
codon
1010 1020 1030 1040 1050
Y P K L E S E F V Y G D Y K V Y D V R K M I A K S E Q E I G K A T A K Y F F Y S N I M N F F K T E I

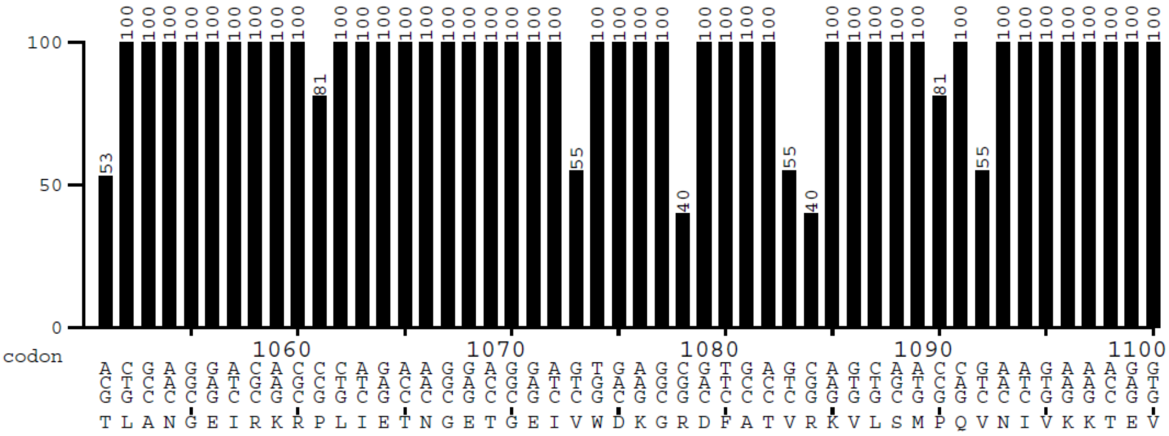
100
50
0
codon
1060 1070 1080 1090 1100
T L A N G E I R K R P L I E T N G E T G E I V W D K G R D F A T V R K V L S M P Q V N I V K K T E V

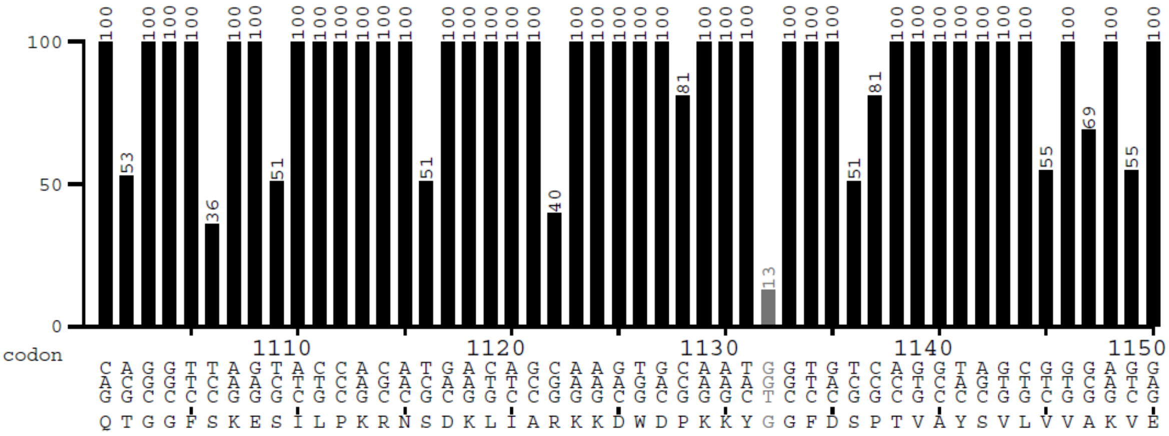
100
50
0
codon
1110 1120 1130 1140 1150
Q T G G F S K E S I L P K R N S D K L I A R K K D W D P K K Y G G F D S P T V A Y S V L V V A K V E

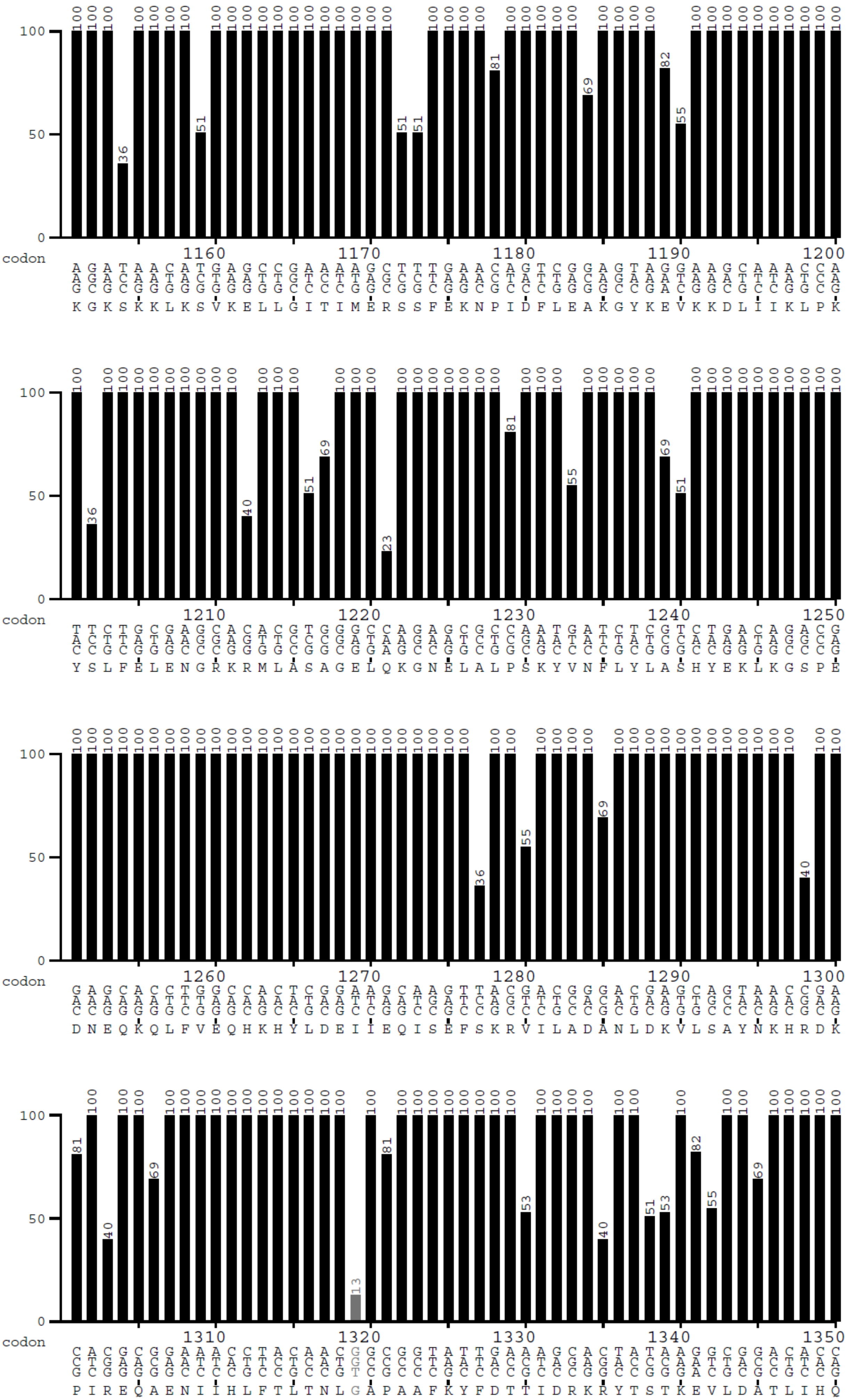
100
50
0
codon
1160 1170 1180 1190 1200
K G K S K K L K S V K E L L G I T I M E R S S F E K N P I D F L E A K G Y K E V K K D L I I K L P K
codon
1210 1220 1230 1240 1250
Y S L F E L E N G R K R M L A S A G E L Q K G N E L A L P S K Y V N F L Y L A S H Y E K L K G S P E
codon
1260 1270 1280 1290 1300
D N E Q K Q L F V E Q H K H Y L D E I I E Q I S E F S K R V I L A D A N L D K V L S A Y N K H R D K
codon
1310 1320 1330 1340 1350
P I R E Q A E N I I H L F T L T N L G A P A A F K Y F D T T I D R K R Y T S T K E V L D A T L I H Q

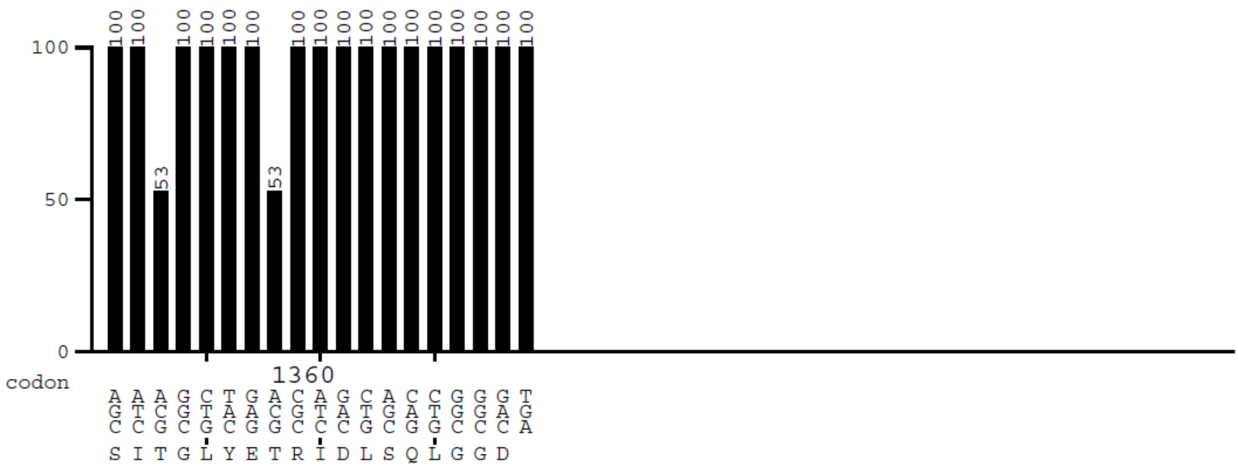


Supplementary Figure 1. *cas9*, optimized for *M. smegmatis* analysed for *D. radiodurans* expression with Graphical Codon Analyzer.

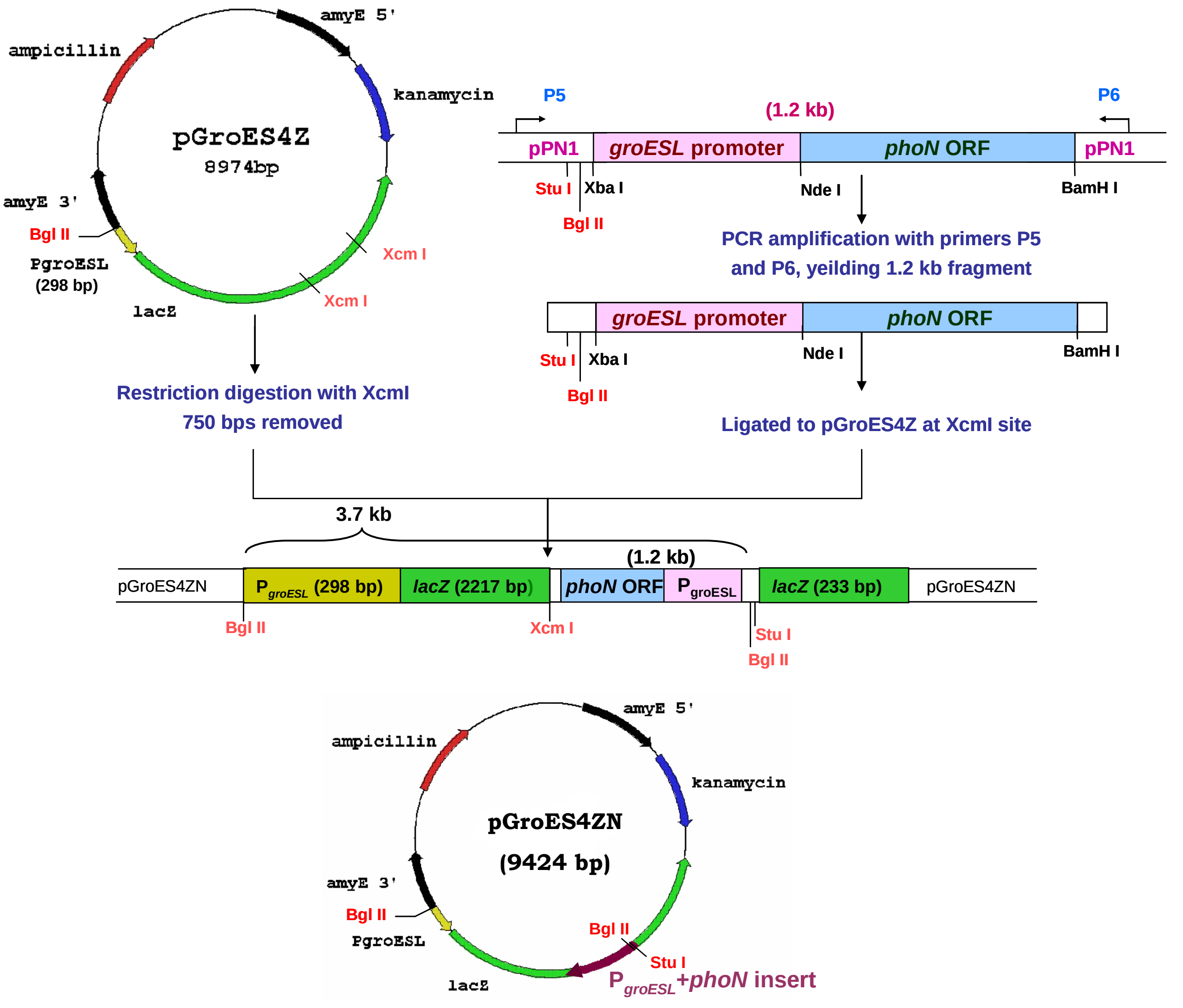


**Supplementary Figure 2:** Cloning of *Salmonella typhi phoN* gene tagged to deinococcal *groESL* promoter into integration vector, pGroES4Z. The deinococcal *groESL* promoter along with the *phoN* ORF was PCR amplified using primers P5 and P6 from plasmid pPN1. The 1.2 kb PCR product was ligated to the XcmI digested pGroES4Z plasmid to yield pGroES4ZN. The relevant restriction enzyme sites are marked in red.

**Supplementary Tables**

**Supplementary Table 1: Plasmids and strains used in study**

| **Strains** | Description | Source/Reference |
|---|---|---|
| *Escherichia coli* JM109 | Cloning strain | Lab strain |
| *Escherichia coli* DH5α | Cloning strain | Lab strain |
| *Deinococcus radiodurans R1* | Wild type | Lab strain |
| *Deinococcus radiodurans R1 phoN+* | *D. radiodurans* with the $P_{groESL}$+*phoN* gene integrated into its genome | This study |
| **Plasmids** | | |
| pRAD1 | *E. coli-D. radiodurans* shuttle vector; $Ap^r$ $Cm^r$; 6.28 kb | Meima et al. 2001 |
| pPN1 | pRAD1 containing *S. enterica* serovar Typhi *phoN* gene with deinococcal *groESL* promoter | Appukuttan et al., 2006 |
| pCRD1 | pRAD1 containing Cas9 under $P_{groESL}$ promoter | This study |
| pCRD2 | pRAD1 containing both Cascade genes and crRNA under $P_{groESL}$ promoter | This study |
| $pCRD2_{PhoN}$ | pCRD2 with crRNA targeting *phoN* gene | This study |
| $pCRD2_{SpORF}$ | pCRD2 with crRNA targeting *ssb* ORF | This study |
| $pCRD2_{SpPRM}$ | pCRD2 with crRNA targeting *ssb* promoter | This study |
| $pCRD2_{PhSb}$ | pCRD2 with crRNA targeting the ORF of both, *phoN* and *ssb*. | This study |
| pSTKT-Cas9 | *E. coli-Mycobacterium* shuttle vector with Cas9 codon optimized for *Mycobacterium* cloned in it | Unpublished results |
| pGroES4Z | Replicative plasmid in *E. coli*, for integrating genes into the *amyE* locus of the chromosome in *D. radodurans*; $Ap^r$, $Kan^r$ | Meima et al., 2006 |
| pGroES4ZN | pGroES4ZN carrying $P_{groESL}$ and *phoN* cloned between the *amyE* flanks for integration into the deinococcal genome. | This study |

**Supplementary Table 2:**

**Codon optimised Sequence of Cascade :**

CAT**atg**
AACCTGCTGATCGACAACTGGATTCCCGTGCGCCCGCGCAACGGTGGCAAGGTGCAGATCATCAAC
CTGCAAAGCCTGTACTGCTCGCGCGACCAGTGGCGCCTGAGCCTGCCCCGCGACGACATGGAGCTG
GCCGCGCTGGCCCTGCTGGTGTGCATCGGCCAGATCATCGCCCCCGCCAAGGACGACGTGGAGTTC
CGCCACCGCATCATGAACCCGCTGACCGAGGACGAGTTCCAGCAGCTGATCGCGCCGTGGATCGAC
ATGTTCTACCTGAACCACGCCGAGCACCCCTTCATGCAGACCAAGGGCGTGAAGGCGAACGACGTG
ACCCCGATGGAAAAGCTGCTGGCCGGCGTGAGCGGCGCCACCAACTGCGCCTTCGTGAACCAGCCT
GGCCAGGGCGAGGCCCTGTGCGGCGGCTGCACCGCCATCGCGCTGTTCAACCAGGCCAACCAGGCC
CCTGGCTTCGGCGGCGGCTTCAAGAGCGGCCTGCGCGGCGGCACCCCGGTGACCACCTTCGTGCGC
GGCATCGACCTGCGCAGCACCGTGCTGCTGAACGTGCTGACCCTGCCCCGCCTCCAGAAGCAGTTCC
CGAACGAGAGCCACACCGAAAACCAGCCCACCTGGATCAAGCCGATCAAGAGCAACGAGAGCATCC
CCGCGAGCAGCATCGGCTTCGTGCGCGGCCTGTTCTGGCAGCCGGCCCACATCGAACTGTGCGACC
CGATCGGCATCGGCAAGTGCAGCTGCTGCGGCCAGGAGAGCAACCTGCGCTACACCGGCTTCCTGA
AGGAAAAGTTCACCTTCACCGTGAACGGCCTGTGGCCCCACCCGCACAGCCCCTGCCTGGTGACCGT
GAAGAAGGGCGAGGTGGAGGAAAAGTTCCTGGCCTTCACCACCAGCGCCCCCAGCTGGACCCAGA
TCAGCCGCGTGGTGGTGGACAAGATCATCCAGAACGAAAACGGCAACCGCGTGGCCGCGGTGGTG
AACCAGTTCCGCAACATCGCGCCCCAGAGCCCGCTGGAGCTGATCATGGGCGGCTACCGCAACAAC
CAGGCCAGCATCCTGGAACGCCGCCACGACGTGCTGATGTTCAACCAGGGCTGGCAGCAGTACGGC
AACGTGATCAACGAGATCGTGACCGTGGGCCTGGGCTACAAGACCGCCCTGCGCAAGGCCCTGTAC
ACCTTCGCGGAAGGCTTCAAGAACAAGGACTTCAAGGGCGCGGGCGTGAGCGTGCACGAAACCGC
CGAGCGCCACTTCTACCGCCAGAGCGAGCTGCTGATCCCCGACGTGCTGGCCAACGTGAACTTCAGC
CAGGCCGACGAGGTGATCGCCGACCTGCGCGACAAGCTGCACCAGCTGTGCGAAATGCTGTTCAAC
CAGAGCGTGGCCCCCTACGCCCACCACCCGAAGCTGATCAGCACCCTGGCCCTGGCGCGCGCCACC
CTGTACAAGCACCTGCGCGAGCTGAAGCCG ca**aggagg** gccCAGCa**at g**gc

**tga**C
GAGATCGACGCGATGGCCCTGTACCGCGCCTGGCAGCAGCTGGACAACGGCAGCTGCGCCCAGATC
CGCCGCGTGAGCGAGCCGGACGAACTGCGCGACATCCCCGCCTTCTACCGCCTGGTGCAGCCCTTC
GGCTGGGAAAACCCGCGCCACCAGCAGGCCCTGCTGCGCATGGTGTTCTGCCTGAGCGCGGGCAA
GAACGTGATCCGCCACCAGGACAAGAAGAGCGAGCAGACCACCGGCATCAGCCTGGGCCGCGCCC
TGGCCAACAGCGGCCGGATCAACGAACGTCGCATCTTCCAGCTGATCCGCGCGGACCGCACCGCGG
ACATGGTGCAGCTGCGCCGCCTGCTGACCCACGCGGAGCCGGTGCTGGACTGGCCCCTGATGGCCC
GCATGCTGACCTGGTGGGGCAAGCGCGAGCGCCAGCAGCTGCTGGAGGACTTCGTGCTGACCACC
AACAAGAACGCG

**taaggaGG**cctt tct

**atg**
AGCAACTTCATCAACATCCACGTGCTGATCAGCCACAGCCCCAGCTGCCTGAACCGCGACGACATGA
ACATGCAGAAGGACGCCATCTTCGGTGGCAAGCGTCGCGTGCGCATCAGCAGCCAGAGCCTGAAGC
GCGCCATGCGCAAGAGCGGCTACTACGCGCAGAACATCGGCGAAAGCAGCCTGCGCACCATCCACC
TGGCCCAGCTGCGCGACGTGCTGCGCCAGAAGCTGGGCGAGCGCTTCGACCAGAAGATCATCGACA

AGACCCTGGCCCTGCTGAGCGGCAAGAGCGTGGACGAGGCGGAAAAGATCAGCGCGGACGCGGT
GACCCCGTGGGTGGTGGGCGAGATCGCCTGGTTCTGCGAACAGGTGGCGAAGGCCGAGGCGGACA
ACCTGGACGACAAGAAGCTGCTGAAGGTGCTGAAAGAGGACATCGCCGCGATCCGCGTGAACCTG
CAACAGGGCGTGGACATCGCCCTGAGCGGCCGGATGGCCACCAGCGGCATGATGACCGAGCTGGG
CAAGGTGGACGGCGCCATGAGCATCGCCCACGCGATCACCACCCACCAGGTGGACAGCGACATCGA
CTGGTTCACCGCCGTGGACGACCTGCAAGAACAGGGCAGCGCCCACCTGGGCACCCAGGAGTTCAG
CAGCGGCGTGTTCTACCGCTACGCCAACATCAACCTGGCCCAGCTCCAGGAAAACCTGGGCGGCGC
CAGCCGCGAGCAGGCCCTGGAAATCGCCACCCACGTGGTGCACATGCTGGCCACCGAGGTGCCCGG
CGCCAAGCAGCGCACCTACGCCGCCTTCAACCCGGCGGACATGGTGATGGTGAACTTCAGCGACAT
GCCCCTGAGCATGGCCAACGCGTTCGAGAAGGCCGTGAAGGCGAAGGACGGCTTCCTCCAGCCGA
GCATCCAGGCCTTCAACCAGTACTGGGACCGCGTGGCCAACGGCTACGGCCTGAACGGCGCTGCCG
CCCAGTTCAGCCTGAGCGACGTGGACCCCATCACCGCCCAGGTGAAGCAGATGCCGACCCTGGAAC
AGCTGAAGAGCTGGGTGCGCAACAACGGCGAGGCG

**Tga**GGAGGCCTT TCT

**atg**
CGCAGCTACCTGATCCTGCGCCTGGCCGGCCCCATGCAGGCGTGGGGCCAGCCGACCTTCGAGGGC
ACCCGCCCCACCGGCCGCTTCCCGACCCGCAGCGGCCTGCTGGGCCTGCTGGGCGCCTGCCTGGGC
ATCCAGCGCGACGACACCAGCAGCCTCCAGGCCCTGAGCGAGAGCGTGCAGTTCGCCGTGCGCTGC
GACGAACTGATCCTGGACGACCGCCGCGTGAGCGTGACCGGCCTGCGCGACTACCACACCGTGCTG
GGCGCCCGCGAGGACTACCGCGGCCTGAAGAGCCACGAAACCATCCAGACCTGGCGCGAATACCTG
TGCGACGCCAGCTTCACCGTGGCCCTGTGGCTGACCCCGCACGCCACGATGGTGATCAGCGAGCTG
GAAAAGGCGGTGCTGAAGCCCCGCTACACCCCGTACCTGGGCCGCCGCAGCTGCCCCCTGACCCAC
CCGCTGTTCCTGGGCACCTGCCAGGCCAGCGACCCGCAGAAGGCGCTGCTGAACTACGAGCCCGTG
GGCGGCGACATCTACAGCGAGGAAAGCGTGACCGGCCACCACCTGAAGTTCACCGCCCGCGACGA
GCCCATGATCACCCTGCCCCGCCAGTTCGCGAGCCGCGAATGGTACGTGATC aa**agg aggC**atgg **at**
**g**tatctcag

**t aa**a
GTGATCATCGCCCGTGCCTGGAGCCGCGACCTGTACCAGCTGCACCAGGGCCTGTGGCACCTGTTCC
CCAACCGCCCCGACGCGGCGCGCGACTTCCTGTTCCACGTGGAGAAGCGCAACACCCCGGAAGGCT
GCCACGTGCTGCTCCAGAGCGCCCAGATGCCGGTGAGCACCGCCGTGGCGACCGTGATCAAGACCA
AGCAGGTGGAGTTCCAGCTCCAGGTGGGCGTGCCCCTGTACTTCCGCCTGCGCGCCAACCCGATCA
AGACCATCCTGGACAACCAGAAGCGCCTGGACAGCAAGGGCAACATCAAGCGCTGCCGCGTGCCCC
TGATCAAGGAAGCCGAACAGATCGCGTGGCTCCAGCGCAAGCTGGGCAACGCGGCGCGCGTGGAG
GACGTGCACCCCATCAGCGAACGCCCGCAGTACTTCAGCGGCGACGGCAAGAGCGGCAAGATCCA
GACCGTGTGCTTCGAAGGCGTGCTGACCATCAACGACGCCCCGGCCCTGATCGACCTGGTGCAGCA
GGGCATCGGCCCTGCCAAGAGCATGGGCTGCGGCCTGCTGAGCCTGGCGCCGCTG **tga**

Red alphabets – Start /stop codons, Highlighted sequence – Shine Dalgarno sequence

**Supplementary Table 3: Oligos used in study**

| oligo Name | 5'-3' Sequence | Description |
|---|---|---|
| Pgrocas-f | CGTCATATGGGGTCCTCCTGTGAGTGAG | For cloning PgroESL into pRAD1 |
| Pgrocas-r | AATCGGATCCCATGTTCAGGGATGGAAGCAC | For cloning PgroESL into pRAD1 |
| CorrPhoNSpa-f | TCGAGAGTTCCCCGCGCCAGCGGGGATAAACCGAAA AGTCGTTATTTACTATTTTT TCTACCACTGAG TTCCCCGCGCCAGCGGGGATAAACCGAGCT | Spacer for targeting PhoN |
| CorrPhoNSpa-r | CGGTTTATCCCCGCTGGCGCGGGGAACTCAGTGGTA GAAAAAAATAGTAAATAACGACTTTTCGGTT TATCCCCGCTGGCGCGGGGAACTC | Spacer for targeting PhoN |
| SpaSsb1f | TCGAGAGTTCCCCGCGCCAGCGGGGATAAACCGTCA TTGACATAATTGACTCTGCTTGTTACTAT GAGTTCCCCGCGCCAGCGGGGATAAACCGAGCT | Spacer for targeting promoter of Ssb |
| SpaSsb1r | CGGTTTATCCCCGCTGGCGCGGGGAACTCATAGTAAC AAGCAGAGTCAATTATGTCAATGA CGGTTTATCCCCGCTGGCGCGGGGAACTC | Spacer for targeting promoter of Ssb |
| SpaSsb3f | TCGAGAGTTCCCCGCGCCAGCGGGGATAAACCGGCC CGAGGCATGAACCACGTCTACCTGATCGG GAGTTCCCCGCGCCAGCGGGGATAAACCGAGCT | Spacer for targeting ORF of Ssb |
| SpaSsb3r | CGGTTTATCCCCGCTGGCGCGGGGAACTCCCGATCAG GTAGACGTGGTTCATGCCTCGGGC CGGTTTATCCCCGCTGGCGCGGGGAACTC | Spacer for targeting ORF of Ssb |
| P5 | GGAGCGGATAACAATTTCACACA | For integration of $P_{groESL}$+*phoN* construct into *Deinococcus* genome |
| P6 | AACGCGGCTGCAAGAATGGTA | For integration of $P_{groESL}$+*phoN* construct into *Deinococcus* genome |
| Amy-1 | CGTATGCCTCACCCTGACATC | Diagnostic primers to confirm *phoN* integration in *D. radiodurans* |
| Amy-2 | AGATTTTGAGACACAACGTG | Diagnostic primers to confirm *phoN* integration in *D. radiodurans* |
| Amy-3 | CTCCTGAGATCTTCCCTGCAG | Diagnostic primers to confirm *phoN* integration in *D. radiodurans* |
| Amy-4 | GATGAGGGAGCAAGTCAGAC | Diagnostic primers to confirm *phoN* integration in *D. radiodurans* |

**Supplementary Table 4: Phosphatase activity in different strains**

| Strain | Plasmid | Phosphatase activity (nmol *p*-NP released/mg protein/min) |
|---|---|---|
| *D. radiodurans* | pRAD1 | 30 ± 5 |
| *D. radiodurans* | pPN1 | 190 ±10 |
| *D. radiodurans* | - | 32 ± 3 |
| *D. radiodurans phoN+* | - | 65 ± 5 |
